# Preoptic BRS3 neurons increase body temperature and heart rate via multiple pathways

**DOI:** 10.1101/2021.03.04.433948

**Authors:** Ramón A. Piñol, Allison S. Mogul, Colleen K. Hadley, Atreyi Saha, Chia Li, Vojtěch Škop, Haley S. Province, Cuiying Xiao, Oksana Gavrilova, Michael J. Krashes, Marc L. Reitman

## Abstract

The preoptic area (POA) is a key region controlling body temperature (Tb), dictating thermogenic, cardiovascular, and behavioral responses to regulate Tb. Known POA neuronal populations reduce Tb when activated; a population that increases Tb upon activation has not yet been reported. We now identify bombesin-like receptor 3 (BRS3)-expressing POA (POA^BRS3^) neurons as having this missing functionality. BRS3 is an orphan receptor that regulates energy and cardiovascular homeostasis, but the relevant neural circuits are incompletely understood. In mice, we demonstrate that POA^BRS3^ neuronal activation increases Tb, heart rate, and blood pressure sympathetically, via projections to the paraventricular nucleus of the hypothalamus and dorsomedial hypothalamus. Acute POA^BRS3^ inhibition reduces Tb. Long-term inactivation of POA^BRS3^ neurons increased Tb variability with exaggerated Tb changes, overshooting both increases and decreases in Tb set point. BRS3 marks preoptic populations that regulate Tb and heart rate, contribute to cold-defense and fine-tune feedback control of Tb. These findings advance understanding of homeothermy, a defining feature of mammalian biology.

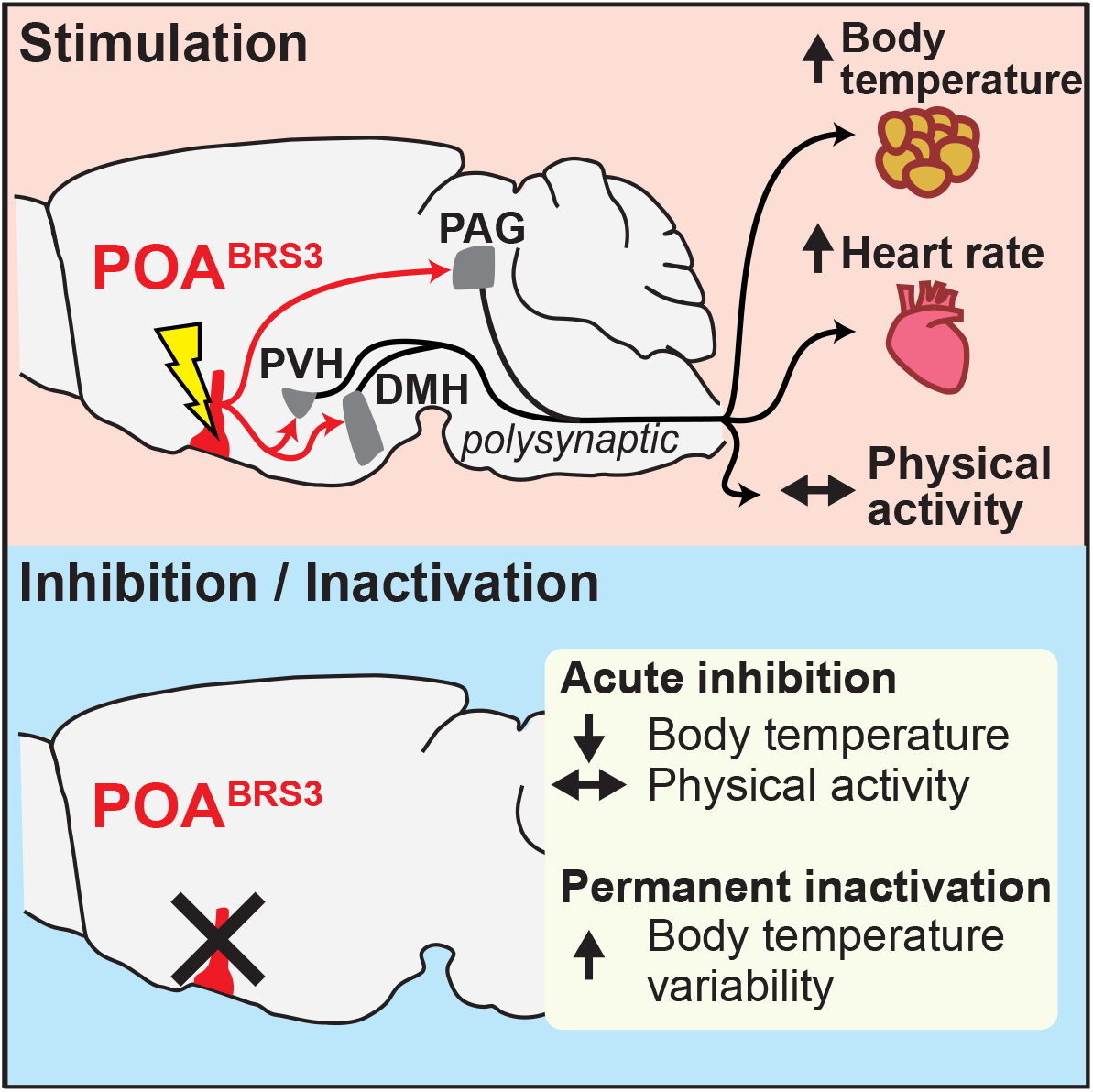
Graphical abstract.

## Introduction

Homeothermy is the property of having a stable core body temperature (Tb), which allows finer control of body processes. Endotherms, including mammals and birds, are homeotherms that use metabolism-generated heat to achieve a warm Tb. To regulate Tb, both environmental temperature (Ta) and Tb must be sensed and evaluated. The organism controls heat generation, including adaptive heat production from brown and beige/brite adipose tissue and muscle. Heat conservation/dissipation is also highly regulated, typically by vasodilation/vasoconstriction and species-determined mechanisms such as panting, sweating, and a variety of behavioral adaptations. The regulation of these processes is orchestrated by the central nervous system. As ultimately survival depends on proper Tb regulation, it is critical to understand this essential physiology.

The preoptic area (POA) is a major integratory hub regulating Tb and cardiovascular responses, and drinking, sleep, parenting, sex, and reward behaviors (Dulac et al., 2014; McHenry et al., 2017; McKinley et al., 2015; Simerly, 1998). The POA contributes to Tb regulation in response to a warm or cold Ta, in producing fever, in hibernation and torpor, and during sleep (Morrison and Nakamura, 2019; Tan and Knight, 2018). Neuron chemotypes in various POA subregions that are activated by warm ambient temperatures and/or reduce Tb when activated include those expressing Vglut2/PACAP/leptin receptor, Vgat, BDNF, galanin, TRPM2, NOS1, QRFP, ERα, and PGDS2 (Abbott and Saper, 2017; Harding et al., 2018; Hrvatin et al., 2020; Kroeger et al., 2018; Moffitt et al., 2018; Song et al., 2016; Takahashi et al., 2020; Tan et al., 2016; Wang et al., 2019; Yu et al., 2016; Zhang et al., 2020; Zhao et al., 2017). Many of these chemotypes specify overlapping populations. POA cold-sensitive neurons drive heat generation and conservation and increase heart rate (HR) (Nakamura and Morrison, 2008). The neuronal identity and circuitry of cold-sensitive neurons is incompletely understood. A proposed mechanism is disinhibition of an inhibitory POA to dorsomedial hypothalamus (DMH) pathway, but an excitatory POA to DMH projection has also been suggested (Dimitrov et al., 2011; Morrison and Nakamura, 2019; da Conceicao et al., 2020). Therefore, a major gap in our understanding of Tb control is that no specific population of POA neurons whose activation increases Tb has been identified. The POA population implicated in fever expresses EP3R and is inhibited by PGE2 to produce fever; this population may not overlap with the cold-defense neurons (Machado et al., 2020). It is not known if POA cold-responsive neurons drive thermogenesis via pathways other than to the DMH.

Bombesin-like receptor 3 (BRS3, BB3, bombesin receptor subtype-3) is an orphan G protein-coupled receptor that regulates energy metabolism and the cardiovascular system (Ohki-Hamazaki et al., 1997). BRS3 is expressed in some peripheral sites (Jensen et al., 2008), but its effects on food intake, metabolic rate, Tb, HR, and blood pressure are due to action in the brain (Guan et al., 2010; Xiao and Reitman, 2016). BRS3 has a restricted brain distribution (Maruyama et al., 2018; Pinol et al., 2018; Zhang et al., 2013), with its metabolic effects attributed to glutamatergic neurons and in part to those expressing MC4R and SIM1 (Xiao et al., 2020; Xiao et al., 2017). Activation of BRS3 neurons in the DMH (DMH^BRS3^) increased energy expenditure, Tb, HR, and blood pressure, while activation of paraventricular nucleus of the hypothalamus (PVH)^BRS3^ neurons reduced food intake (Pinol et al., 2018). We now demonstrate that POA^BRS3^ neurons actively contribute to cold-defense and to the feedback control of Tb.

## Results

### Optogenetic activation of POA^BRS3^ neurons rapidly increases Tb, heart rate, and blood pressure

Neurons expressing BRS3 in the preoptic area (POA^BRS3^) were activated by exposure to a cold ambient temperature, as demonstrated by increased Fos expression (Figure 1a,b) (Pinol et al., 2018), so we tested the ability of POA^BRS3^ neurons to control Tb, HR, and mean arterial pressure (MAP). Optogenetic stimulation of POA^BRS3^ neurons increased Tb by 1.2 ± 0.2 °C, HR by 134 ± 18 bpm, and MAP by 20.4 ± 1.4 mm Hg, with no increase in physical activity and no changes in control mice (Figure 1c,d). We varied stimulation times to characterize the onset of the HR and MAP increases. Stimulation for 0.5 s increased HR and MAP detectably. The half-maximal HR response was elicited with ∼2 s of stimulation, with slightly more time needed for MAP. Stimulation for 20 s maximally increased both HR and MAP (Figure 1e,f). The Tb response was slow (50% at 6.3 ± 0.5 min) due to the body’s heat capacity. Thus, activation of POA^BRS3^ neurons increases Tb, HR, and MAP independent of physical activity.

**Figure 1.**
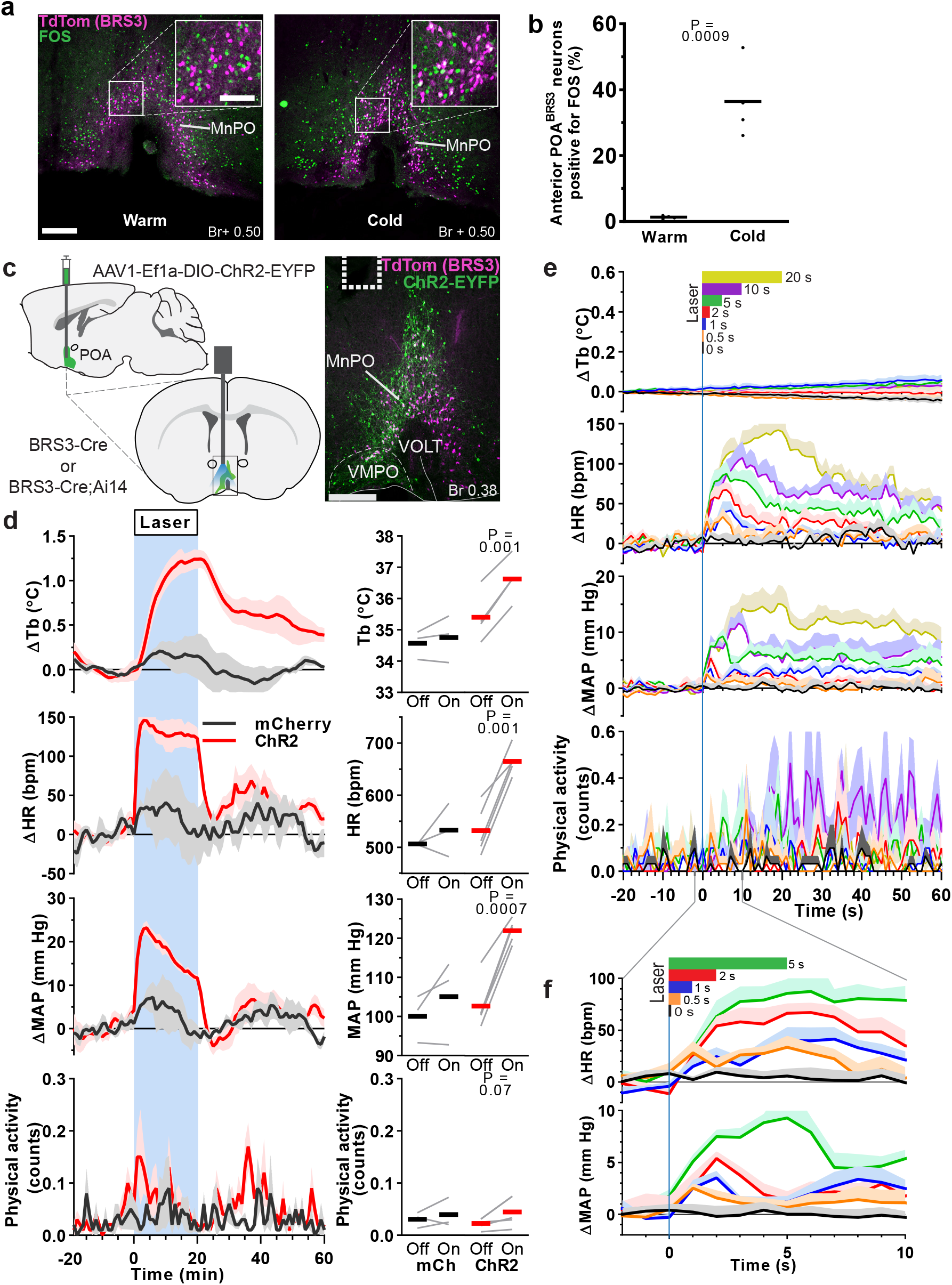
POA^BRS3^ neurons are cold-sensitive and optogenetic stimulation increases body temperature, heart rate and mean arterial pressure. a) FOS expression (green) in BRS3 neurons (magenta) of 4 h cold-(4 °C) or warm-exposed (30 °C) mice. Scale bar is 200 µm, 80 µm in inset. b) Average and individual percentages of colocalization of FOS in anterior (Bregma +0.50 and +0.38) POA^BRS3^ neurons (n = 4/group). P values from unpaired t test, cold vs warm. Data are re-graphed from (Pinol et al., 2018). c) Schematic of virus injection (top left) and placement of optic fiber (bottom left). ChR2-EYFP expression (green) in BRS3 (magenta) neurons in the MnPO of a BRS3-Cre;Ai14 mouse (right). Dotted line indicates fiber placement. Scale bar is 200 µm. MnPO – median preoptic area; VMPO – ventromedial preoptic area; VOLT – vascular organ of lamina terminalis. d) Tb, HR, MAP, and physical activity response to 20 min laser stimulation (blue interval; 3s on 1s off; 20 Hz; 10 ms pulses). mCherry controls (black, n = 3), ChR2 (red, n = 5). Data are average of 5 epochs/mouse, relative to epoch baseline (−20 to −1 min); mean ± s.e.m. Quantitation in right panels (intervals: Off, −10 to −1 min; On, 10 to 19 min for Tb and 0 to 9 min for HR, MAP, and physical activity; bars, means; gray lines, individual animals; P values from paired t test, Off vs On). e, f) Brief optogenetic stimulation increases HR and MAP, but not Tb or physical activity. Laser (continuous pulses; 20 Hz; 10 ms pulses) stimulation is indicated at top in color-coded bars. Data are mean + s.e.m., n=5 mice (average of 10 epochs/mouse), relative to epoch baseline (−59 to 0 s).

The Tb increase caused by POA^BRS3^ neuron activation is in the opposite direction of that observed in most other POA neuron populations, activation of which reduces Tb (Morrison and Nakamura, 2019). This makes POA^BRS3^ neurons functionally distinct from other POA populations. We next used a Cre-off ChR2-expressing virus to selectively activate POA neurons that do not express BRS3 (POA^nonBRS3^). We compared non-selective stimulation of POA neurons (POA^All^) and stimulation of POA^nonBRS3^ neurons with stimulation of POA^BRS3^ neurons (Figure 2). Optogenetic activation of either POA^All^ or POA ^nonBRS3^ neurons drastically decreased Tb by almost 3 °C at the end of the 20 min stimulation, with no sign of plateauing. Stimulation of either POA^All^ or POA^nonBRS3^ neurons also massively increased physical activity, which is opposite of the usual behavior during a Tb decrease. Together, these data demonstrate that POA^BRS3^ neurons are a distinct population of Tb- and HR-regulating neurons that function in the opposite direction from previously described POA neuronal populations.

**Figure 2.**
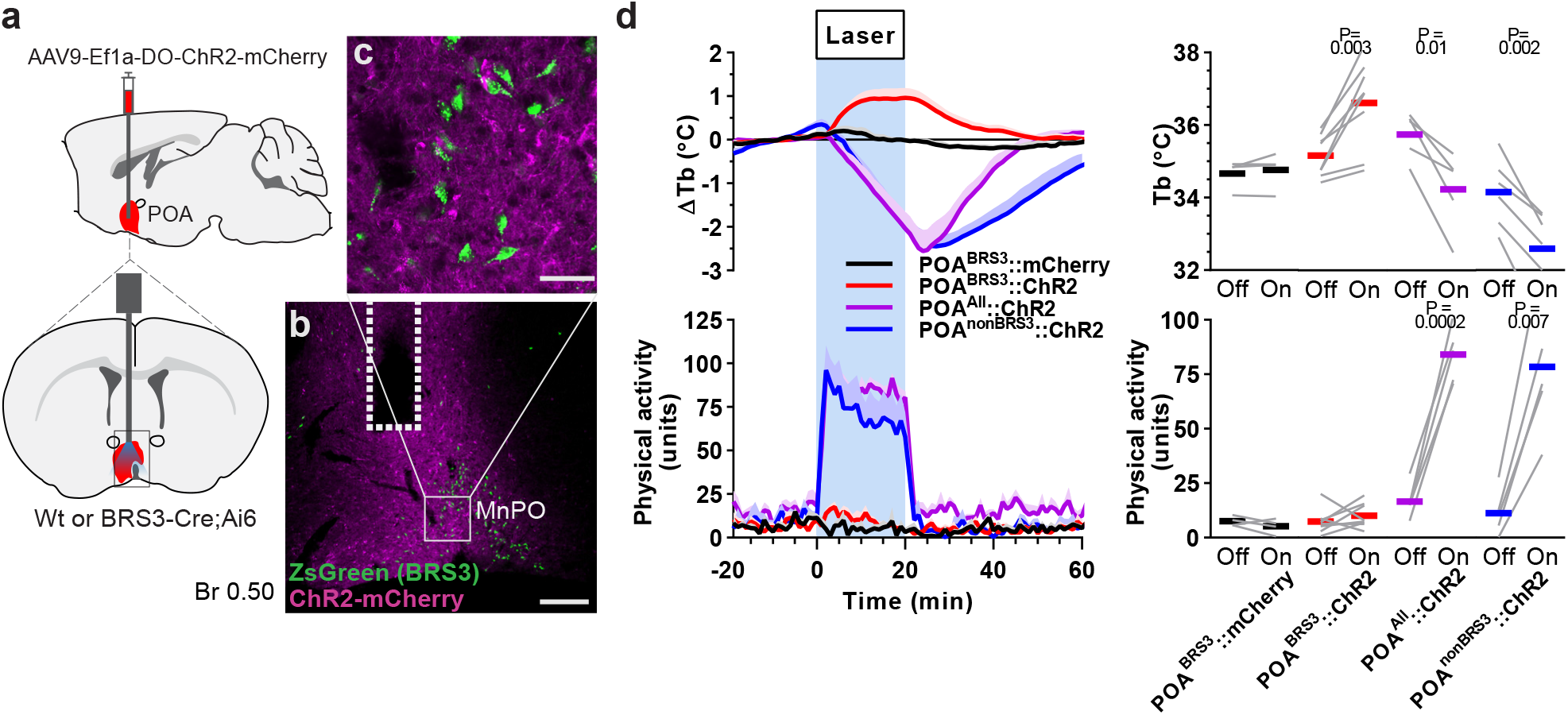
Global stimulation of POA neurons decreases body temperature and increases physical activity. a) Schematic of virus injection (top left) and optic fiber placement (bottom left). b,c) ChR2-EYFP expression in the MnPO of a BRS3-Cre;Ai14 mouse, with detail showing absence of ChR2-mCherry (magenta) expression in BRS3 (green) neurons. Dotted line indicates fiber placement. Scale bar: b) 200 µm, c) 40 µm. d) Tb and physical activity response to 20 minute laser stimulation (blue interval; 3s on 1s off; 20 Hz; 10 ms pulses). POA^BRS3^::mCherry (black, n=4), POA^BRS3^::ChR2 (red, n=8); POA^All^::ChR2 (magenta, n = 5), POA^nonBRS3^::ChR2 (blue, n = 6). Data are average of 5 epochs/mouse, relative to epoch baseline (−20 to −1 min); mean + s.e.m. Quantitation in right panels (intervals: Off, −10 to −1 min; On, 10 to 19 min for Tb and 0 to 9 min for physical activity; bars, means; gray lines, individual animals; P values from paired t test, Off vs On).

### Inhibition of POA^BRS3^ neurons reduces Tb and cold defense

We used chemogenetics to manipulate POA^BRS3^ neurons bidirectionally. Stimulation of the activating DREADD, hM3Dq, with CNO had two effects (Figure 3a,b). First, the initial, handling-associated Tb increase returned to baseline more quickly after CNO than vehicle (CNO, 53 ± 4 min vs vehicle, 71 ± 4 min; p = 0.03), with no difference in physical activity between treatments. Then, starting about 120 minutes after dosing, the Tb in the CNO-treated mice increased by 0.72 ± 0.09 °C, with no increase after vehicle treatment (−0.08 ± 0.07 °C; p = 0.00002 CNO vs vehicle). Thus, chemogenetic stimulation of POA^BRS3^ neurons produces a biphasic response, initially lowering Tb, and then raising Tb. The delayed Tb increase after chemogenetic activation contrasts with the near-immediate onset of the Tb increase upon optogenetic stimulation. The biphasic response suggests that there may be more than one population of POA^BRS3^ neurons regulating Tb.

**Figure 3.**
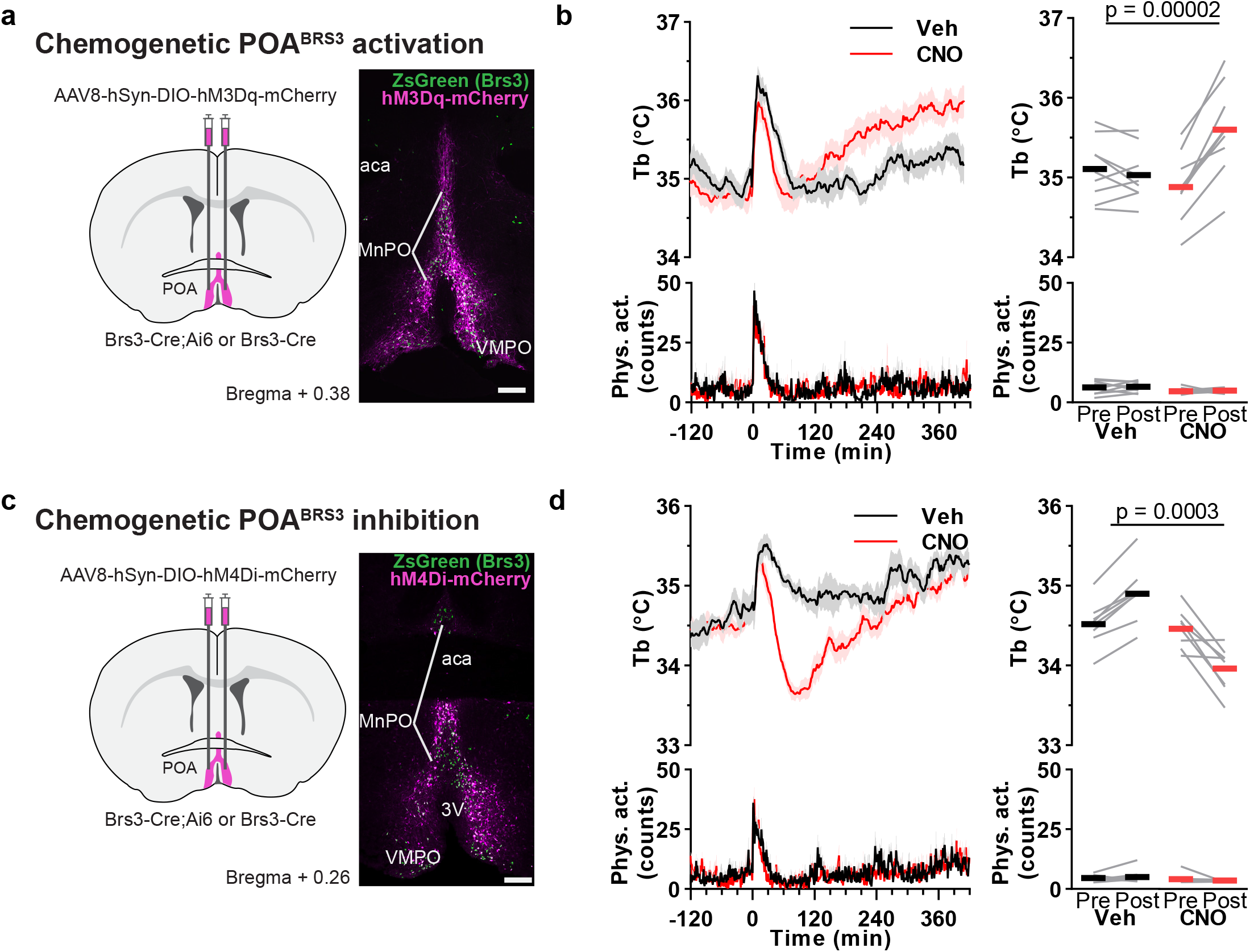
Bi-directional chemogenetic manipulation of POA^BRS3^ neurons. a,c) Schematic of virus injection and example of hM3Dq-mCherry (a) or hM4Di-mCherry (c) expression (magenta) in BRS3 (green) neurons in the anterior POA of a BRS3-Cre;Ai14 mouse. Scale bar is 200 µm. b,d) Tb and physical activity response to CNO (1 mg/kg) or vehicle in POA^BRS3^::hM3Dq mice (c; n = 9, mean of two or three trials; Pre: −150 to −30; Post: 180 to 300 min) or POA^BRS3^::hM4Di (d; n = 8, mean of three trials; Pre: −150 to −30; Post: 60 to 180 min) mice. P values from paired, two-sided t-test on change from baseline, CNO vs vehicle. Data are mean + s.e.m in bottom left panel and ± s.e.m. in top left panel. Bars, means; gray lines, individual animals in right panels. See also Figure S1.

Chemogenetic inhibition of POA^BRS3^ neurons expressing the hM4Di DREADD decreased Tb by 0.50 ± 0.12 °C (vs an increase of 0.39 ± 0.05 °C with vehicle; p = 0.0003 CNO vs vehicle; Figure 3c,d). The Tb reduction started within minutes, blunting the handling-induced Tb increase with no effect on physical activity. These results demonstrate that POA^BRS3^ neuronal activity maintains Tb in a 22 °C environment and suggests that POA^BRS3^ neurons contribute to cold defense.

### Chemogenetic stimulation of BNST^BRS3^ neurons does not change Tb, physical activity or food intake

The bed nucleus of the stria terminalis (BNST) can regulate Tb (Craig, 2018; Schneeberger et al., 2019) and food intake (Jennings et al., 2013) and contains BRS3-expressing neurons (BNST^BRS3^). Chemogenetic activation of BNST^BRS3^ neurons expressing an excitatory DREADD had no effect on Tb, physical activity, or food intake (either reduction or increase) (Figure S1). Thus, BNST^BRS3^ neurons do not appear to control Tb and BNST^BRS3^ and POA^BRS3^ neurons have distinct functions.

### Optogenetic stimulation of POA^BRS3^ neuron projections to PVH, DMH or PAG increases Tb

To identify targets of POA^BRS3^ neurons, we used viral anterograde tracing (Figure S2a-c). POA^BRS3^ neurons had dense local projections to preoptic regions and major projection fields in the PVH, DMH, and raphe pallidus (RPa). Moderate levels of projections were to the periaqueductal grey (PAG) and paraventricular nucleus of the thalamus (PVT). Other areas containing fiber terminals were the lateral septum, lateral hypothalamus, supraoptic nucleus, locus coeruleus, and Barrington’s nucleus.

We next examined which POA^BRS3^ neuron projections increase Tb. Cre-dependent ChR2-expressing virus was injected into the POA of BRS3-Cre mice and an optic fiber was implanted over the PVH, DMH, PVT, PAG, or RPa. Optogenetic stimulation of POA^BRS3^→PVH, POA^BRS3^→DMH, or POA^BRS3^→PAG axons increased Tb by 1.2 ± 0.2 °C, 1.0 ± 0.2 °C, or 0.6 ± 0.05 °C, respectively (Figure 4a-c). Optogenetic stimulation of POA^BRS3^→PVT (Figure 4d) and POA^BRS3^→RPa (Figure 4e) axons did not change Tb. It is possible that PVH stimulation could activate POA^BRS3^→DMH fibers passing through the PVH, although this would not be expected to generate the observed full response. The projection-specific differences suggest that the downstream pathways of POA^BRS3^ neurons differentially contribute to Tb regulation. Laser light delivery in mCherry-expressing control mice did not alter Tb. Physical activity was slightly increased during activation of POA^BRS3^→PVH mice, but not in the other areas. These experiments show that POA^BRS3^ neurons project to multiple brain nuclei, and stimulation of three of these populations increases Tb.

**Figure 4.**
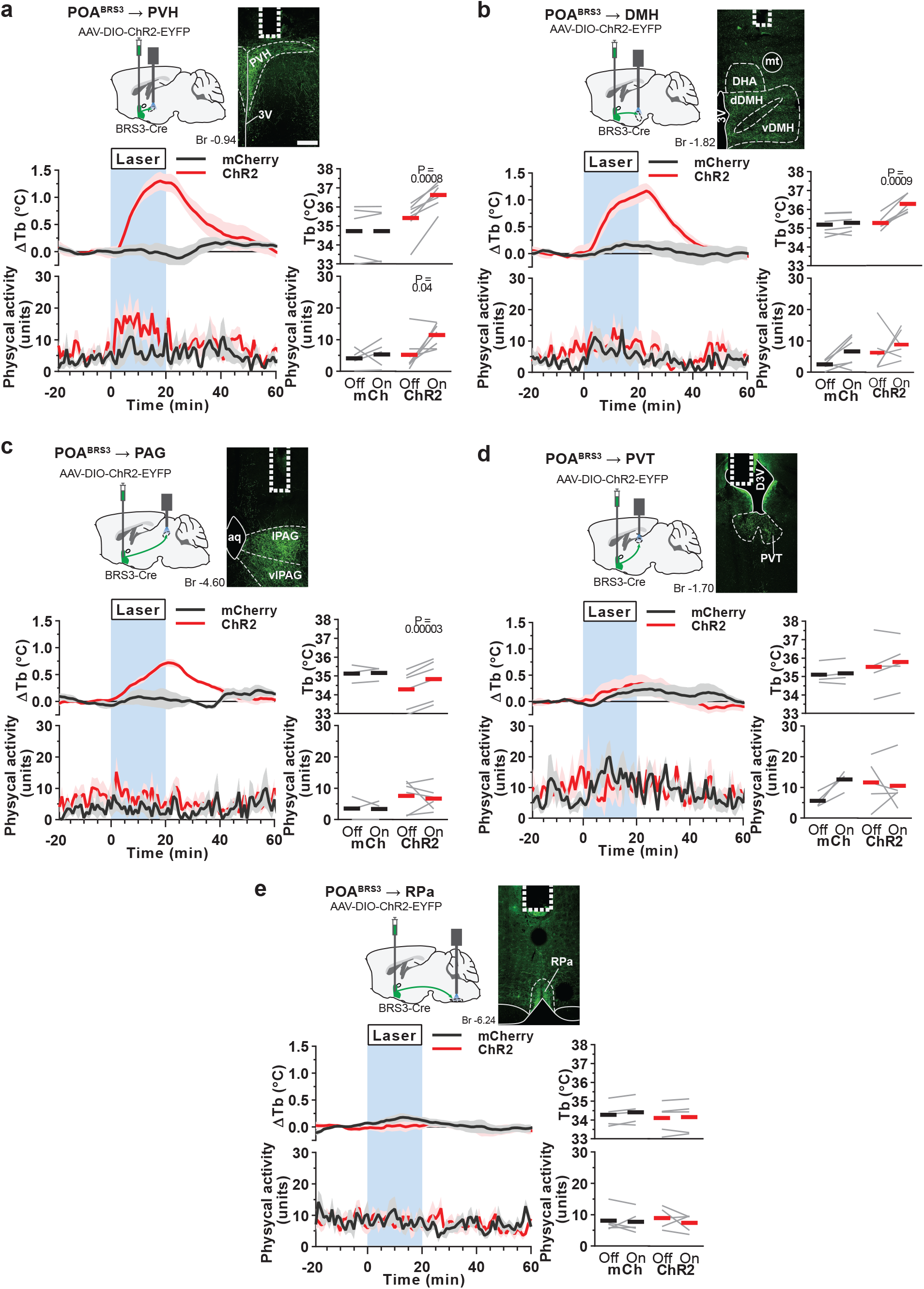
Optogenetic stimulation of POA^BRS3^→PVH, POA^BRS3^→DMH, or POA^BRS3^→PAG axons increases Tb. a-e) Top left: schematic of virus injection and placement of optic fiber. Top right: POA^BRS3^ projections expressing ChR2-EYFP (green) and fiber placement (dotted line). Scale bar is 200 µm. Bottom left: Tb and physical activity response to 20 min laser stimulation (blue interval; 3s on 1s off; 20 Hz; 10 ms pulses). mCherry controls (black, n = 4-5), ChR2 (red, n = 5-7). Data are average of 5-10 epochs/mouse, relative to epoch baseline (−20 to −1 min); mean ± s.e.m. Quantitation in bottom right panels (intervals: Off, −10 to −1 min; On, 10 to 19 min for Tb and 0 to 9 min for physical activity; bars, means; gray lines, individual animals; P values from paired t test, Off vs On). 3V - third ventricle; Aq – aqueduct; DHA – dorsal hypothalamic area dDMH - dorsal part of the dorsomedial hypothalamus; D3V – dorsal part of third ventricle; PVH – paraventricular nucleus of the hypothalamus; PVT – paraventricular nucleus of the thalamus; lPAG - lateral periaqueductal grey; vDMH – ventral part of the dorsomedial hypothalamus; vlPAG – ventrolateral periaqueductal grey; PVH – paraventricular nucleus of the hypothalamus; RPa – raphe pallidus. See also Figures S2, S3.

The POA to DMH pathway is a previously characterized pathway for Tb regulation (Morrison and Nakamura, 2019). Therefore, we assessed if POA^BRS3^→DMH neurons have collaterals. We found that POA^BRS3^→DMH neurons have collaterals to PVH, PVT, PAG, and RPa (Figure S2d-f). Other areas in which we observed collateral fibers strongly overlapped with the anterograde tracing experiment and included the lateral hypothalamus, supraoptic nucleus, locus coeruleus, and Barrington’s nucleus.

### POA^BRS3^→PVH and POA^BRS3^→DMH neurons activate brown adipose tissue

To determine if the Tb increase is mediated by retaining heat through tail vasoconstriction and/or by brown adipose tissue (BAT) activation, we measured skin temperature using infrared imaging (Figure S3a). Tail temperature (T_tail_) was unchanged during the optogenetic stimulation (Figure S3b), indicating a lack of vasodilation, as the tail was already vasoconstricted at 25 °C.

We used temperatures of the interscapular (T_BAT_) and lumbar (T_lumbar_) regions as biomarkers of BAT temperature and Tb, respectively, although one cannot rule out unmeasured localized differential changes in local air temperature or skin blood flow with this assay. T_BAT_ is ∼1 °C warmer than T_lumbar_, consistent with heat production by BAT with transfer to the body. Optogenetic stimulation of POA^BRS3^→PVH or POA^BRS3^→DMH terminals each increased T_BAT_ and T_lumbar_ (Figure S3c,d). The T_BAT_-T_lumbar_ difference increased at the onset of stimulation only in the POA^BRS3^→DMH group and the gradient was maintained during warming, indicating continued BAT activation. Taken together, the results suggest that both POA^BRS3^→PVH and POA^BRS3^→DMH projections activate BAT.

### POA^BRS3^→PVH and POA^BRS3^→DMH neurons stimulate heart rate through the sympathetic nervous system

Since activation of POA^BRS3^ neurons increased HR and MAP, we queried whether this occurred via projections to the PVH or DMH. Optogenetic stimulation of either POA^BRS3^→PVH or POA^BRS3^→DMH terminals increased HR (by 94 ± 9 bpm or 89 ± 27 bpm, respectively) and MAP (by 10.7 ± 2.7 mm Hg or 10.1 ± 3.4 mm Hg, respectively) (Figure 5a). HR and MAP did not change in the mCherry controls and physical activity did not increase significantly in any group.

**Figure 5.**
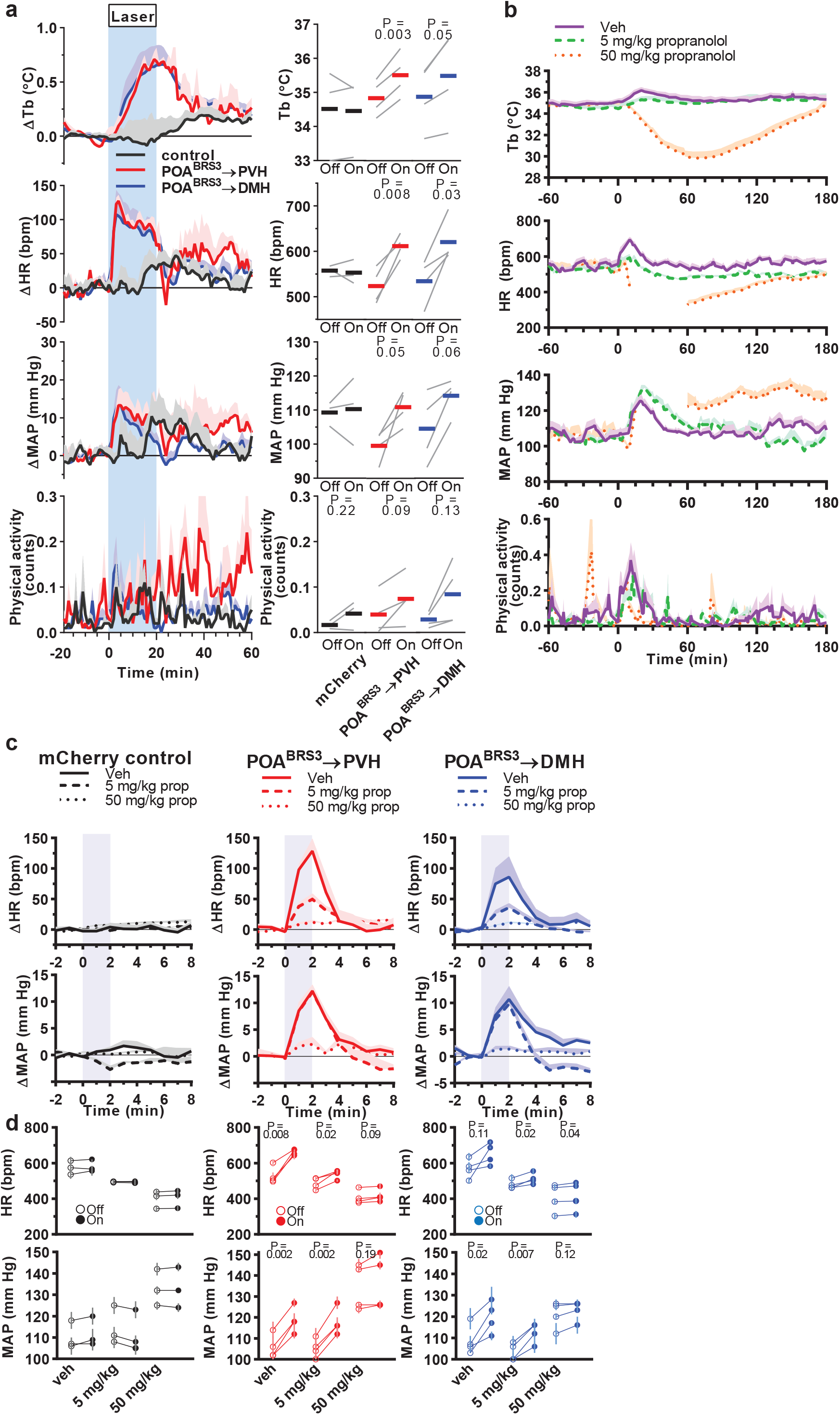
POA^BRS3^→PVH and POA^BRS3^→DMH neurons increase Tb and HR via the sympathetic nervous system. **a**) Tb, HR, MAP, and physical activity response to 20 min laser stimulation (blue interval; 3s on 1s off; 20 Hz; 10 ms pulses). mCherry controls (black, n = 3), POA^BRS3^→PVH (red, n = 4), POA^BRS3^→DMH (blue, n = 4); gray lines, individual animals. Data are average of 5 epochs/mouse, relative to epoch baseline (−20 to −1 min); mean ± s.e.m. Quantitation in bottom right panels (intervals: Off, −10 to −1 min; On, 10 to 19 min for Tb and 0 to 9 min for HR, MAP, and physical activity; P values from paired t test, Off vs On). In **b-d**, propranolol (vehicle, 5, or 50 mg/kg i.p., as indicated) was injected at time 0. Starting at 30 min, consecutive epochs of 2-min laser stimulation (1s on 1s off; 20 Hz; 10 ms pulses) and 8 min laser off were performed. **b**) Effect of propranolol on Tb, HR, MAP, and physical activity. To isolate the effects of propranolol without confounding by optogenetic stimulation, minutes 6-10 (from laser onset) of each 10-minute epoch are graphed. Time 0 is vehicle/propranolol injection. Data are mean + s.e.m. n=11 mice, pooled data. In the 50 mg/kg group, mean is not graphed between 15-60 min as blood pressure signal was lost in most mice; all mice have MAP and HR data after 60 min. Responses to drug (during 75-155 minutes) were compared to baseline (−60 to 0 minutes) and tested with one-way ANOVA, Tukey’s multiple comparisons test (5 mg/kg: HR, p = 0.03 and MAP, p = 0.45; 50 mg/kg: HR, p < 0.0001). c,d) Effect of stimulating POA^BRS3^→PVH or POA^BRS3^→DMH neurons on HR and MAP during propranolol treatment. mCherry controls (black, n = 3), POA^BRS3^→PVH (red, n = 4), POA^BRS3^→DMH (blue, n = 4). Data are mean + s.e.m. of 8 epochs (during 75-155 min). In c, data in each epoch was normalized to its baseline (−1 to −0 min) and the effect of laser stimulation (blue) is depicted as change from baseline. Data are mean + s.e.m. In d, the individual mouse data are presented. Open symbols are baseline (Off, from −1 to 0 minutes) and closed symbols are stimulated (On, from 1 to 2 minutes). P values from paired t test, Off vs On. Data are mean ± s.e.m. of 8 epochs (during 75-155 min). See also Figure S4.

A rapid increase in HR could be due to activation of the sympathetic and/or inhibition of the parasympathetic nervous system. We used propranolol, a β-adrenergic antagonist (blocking β1 and β2 better than β3), to inhibit the sympathetic nervous system. Propranolol’s ED_50_ for Tb reduction was 48 mg/kg i.p. in wild-type mice (Figure S4). Propranolol at 5 mg/kg reduced HR (−68 ± 12 bpm, p = 0.03) but did not change MAP (−4.4 ± 2.9 mm Hg, p = 0.45). The 50 mg/kg dose profoundly reduced HR (−133± 23 bpm, p < 0.0001) and had a biphasic effect on MAP, first reducing it to levels unreliably undetectable by the intra-aortic telemetry, then increasing it (Figure 5b). The effect of propranolol on the optogenetic responses of the POA^BRS3^→PVH and POA^BRS3^→DMH mice were similar: the HR increase was inhibited by 61 ± 5 % and 46 ± 12 %, respectively, with 5 mg/kg and 91 ± 2 % and 82 ± 7 %, respectively, with 50 mg/kg (Figure 5c,d). Propranolol’s effects on the MAP increase were also similar in the POA^BRS3^→PVH and POA^BRS3^→DMH mice, but the MAP and HR dose responses differed. The MAP increase was not inhibited by 5 mg/kg, but was inhibited 80 ± 12 % and 84 ± 7 %, respectively, with 50 mg/kg (Figure 5c,d). Tb and physical activity were not changed by this brief optogenetic stimulation. The biphasic MAP response to 50 mg/kg propranolol complicates interpretation of the optogenetic effects. In contrast, the 5 mg/kg propranolol dose robustly suppressed optogenetic-driven HR increases in POA^BRS3^→PVH and POA^BRS3^→DMH mice, suggesting that these are sympathetic effects. The 5 mg/kg dose of propranolol did not affect MAP, suggesting divergence in the regulation of HR and MAP by these POA^BRS3^ neurons. Taken together, the results indicate that POA^BRS3^→PVH and POA^BRS3^→DMH projections increase HR through the sympathetic nervous system.

### POA^BRS3^→PVH and POA^BRS3^→DMH neurons are mixed glutamatergic and GABAergic populations

We next assessed the neurotransmitters used by POA^BRS3^ neurons. In the median preoptic area (MnPO), 46 ± 7% of BRS3 neurons were positive for Gad2 (Figure S5a,b). Other preoptic BRS3-expressing regions, ventromedial preoptic area (VMPO), ventrolateral preoptic area (VLPO), and medial preoptic area (MPA), had 56 ± 7%, 54 ± 12%, and 40 ± 5% colocalization, respectively. While Gad2 is often used as a marker for GABAergic neurons, it is also expressed in some preoptic glutamatergic (Vglut2-expressing) neurons. However, there are no identified BRS3 populations that express both Vglut2 and Vgat in the POA (Moffitt et al., 2018). The absence of Gad2 expression in a fraction of the preoptic BRS3 neurons suggests that a sub-population of POA^BRS3^ neurons is glutamatergic.

We used ChR2-assisted circuit mapping to test if POA^BRS3^→PVH and POA^BRS3^→DMH projections use glutamate or GABA (Figure 6a). In PVH- or DMH-containing brain slices, optogenetic stimulation of POA^BRS3^ axons elicited both excitatory and inhibitory post-synaptic currents (Figure 6b,c). Thus, both the POA^BRS3^→PVH and POA^BRS3^→DMH neuron populations consist of both glutamatergic and GABAergic subpopulations.

**Figure 6.**
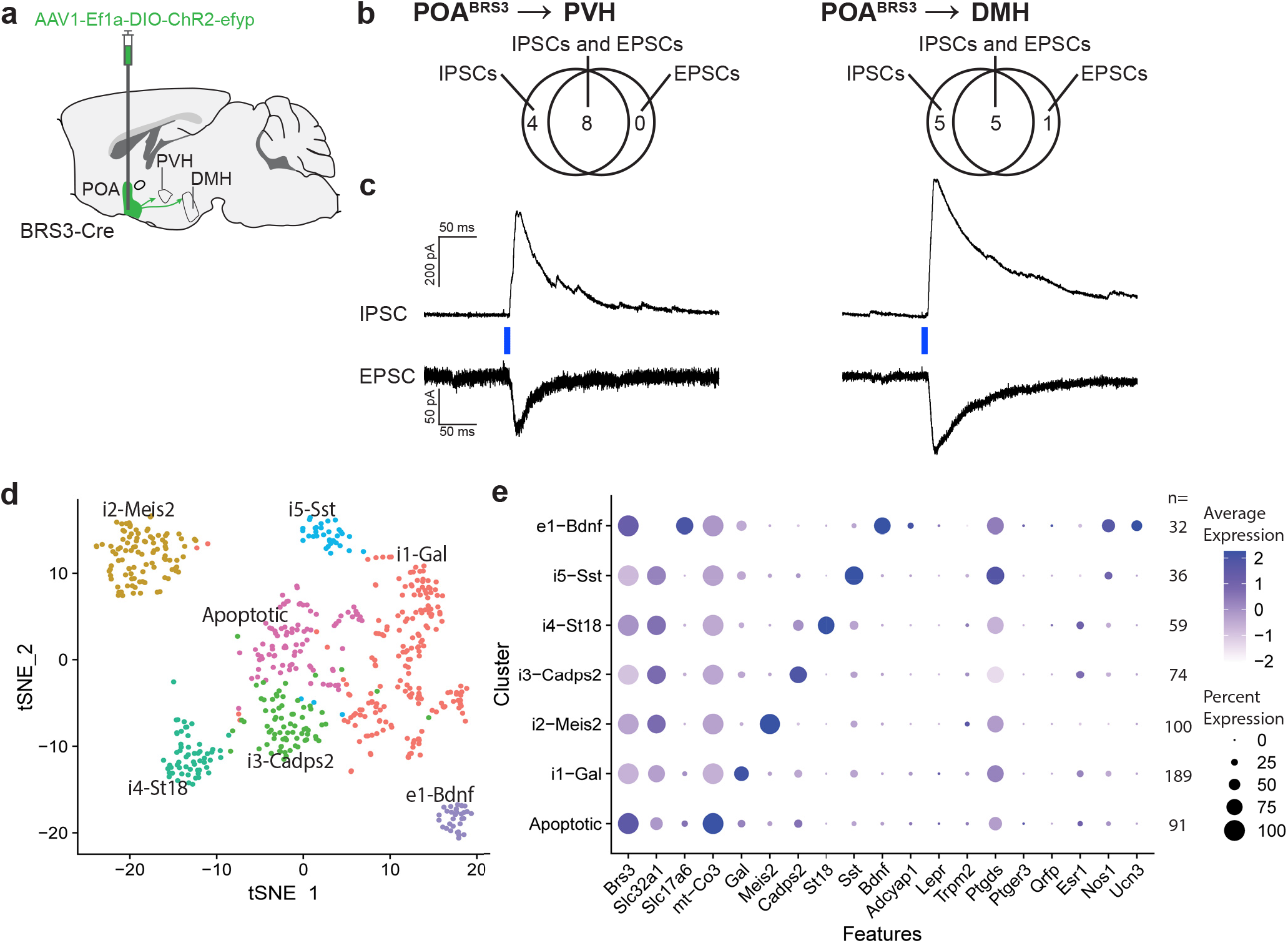
POA^BRS3^→PVH and POA^BRS3^→DMH neurons are both inhibitory and excitatory and POA^BRS3^ are in multiple clusters. a) Schematic showing injection of Cre-dependent ChR2-EYFP-expressing AAV in the POA and projections to PVH and DMH. b) Venn diagrams showing the number of POA^BRS3^→PVH and POA^BRS3^→DMH neurons with each type of postsynaptic current. EPSC, excitatory postsynaptic current; IPSC, inhibitory postsynaptic current. c) Voltage-clamp trace of POA^BRS3^→PVH and POA^BRS3^→DMH stimulation in PVH- and DMH-containing brain slices. Recordings were made in the presence of tetrodotoxin (500 nM) and 4-aminorpyridine (100 µM), showing monosynaptic inhibitory and excitatory input from the POA to both downstream hypothalamic areas. IPSCs recorded at +10 mV and EPSCs at –55 mV. Traces are mean of 10 – 15 stimulations per cell. d) tSNE plot of BRS3 neuron mRNA expression in the POA region with data from (Moffitt et al., 2018). e) Expression of selected mRNAs in the POA region BRS3 clusters. See also Figure S5.

Single-cell RNA sequencing data show that BRS3 is expressed in multiple POA region neuron populations (Moffitt et al., 2018). Reanalysis limited to the 581 BRS3-positive neurons in this dataset (avoiding false negatives due to low BRS3 expression) indicates that there are at least one glutamatergic and five GABAergic populations (Figures 6d,e, S5c). The e1 glutamatergic cluster may contain multiple subclusters, as the cluster heterogeneously expresses markers found in thermoregulatory neurons (eg., Bdnf, Adcyap1, Ptgds, and Nos1) and 44% are positive for galanin. Two clusters, i2-Meis2 and i5-Sst may be in regions neighboring the POA, based on the expression pattern of some of the marker mRNAs (Figure S5d). These results demonstrate that there are multiple populations of POA^BRS3^ neurons.

### Permanent silencing of POA^BRS^ neurons increases Tb variability, exaggerates Tb changes and delays adaptation to Ta below thermoneutrality

To study the effect of chronic inactivation of POA^BRS3^ neurons, we used Cre-dependent tetanus toxin (TeNT)-expressing virus (POA^BRS3^::TeNT mice; control mice expressed EYFP) (Figure 7a,b, S6a). After inactivation, the POA^BRS3^::TeNT mice gained less weight than control mice, with decreased food intake during weeks 4-6 after starting tamoxifen treatment (Figure S6b,c).

**Figure 7.**
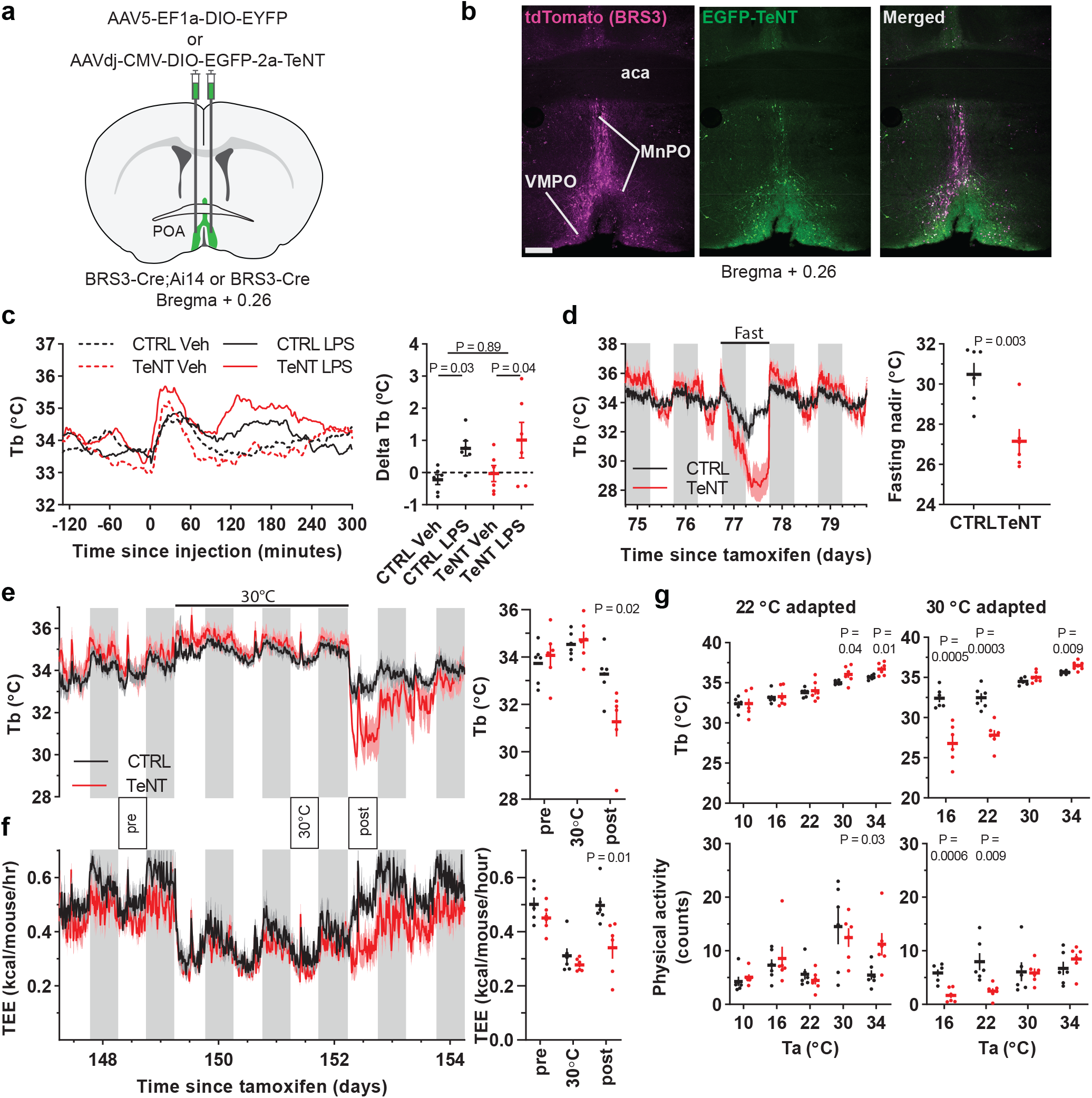
Silencing POA^BRS3^ neurons increases Tb variability and exaggerates Tb changes. a) Schematic of virus injection into the POA of BRS3-Cre mice (control, AAV-DIO-EYFP; silencing, AAV-DIO-EGFP-TeNT). b) Images of a POA^BRS3^;Ai14::TeNT mouse (BRS3, magenta; TeNT, green). aca – anterior commissure; MnPO – median preoptic area; VMPO – ventromedial preoptic area c) Tb response to lipopolysaccharide (LPS, 100 µg/kg, i.p.) or vehicle (saline) and ΔTb (Tb_120to210_ minus Tb_-120to-30_). Data are mean ± s.e.m. (s.e.m. omitted from left for visual clarity); P value, paired t test between vehicle and LPS and unpaired t test with unequal variance between delta Tb (LPS minus Veh) of CTRL and TeNT groups. d) Tb response to 24 h food deprivation and fasting Tb nadir (mean ± s.e.m). P value, unpaired t test. e,f) Tb and total energy expenditure (TEE) of mice at 22 °C, then 3 days at 30 °C, then 22 °C. Quantitation in right panels of the indicated light phase12-h intervals; P value, unpaired t test. g) Acute response to various Ta in mice acclimated to 22 °C (left) or 30 °C (right). After >5 day of acclimation, during light phase mice were exposed to 180 min of the indicated Ta. The mean Tb and physical activity at 60-180 min is shown. P value, unpaired t test. In all panels, n=6 mice/group. See also Figure S6.

The baseline Tb phenotype of the POA^BRS3^::TeNT mice was subtle, consisting of unchanged 24-h mean Tb, generally non-significantly higher dark and lower light phase mean Tb, and a larger SD for both of these. More sensitive metrics were the circadian amplitude (Tb_dark_-Tb_light_) and the Tb span (95^th^ – 5^th^ Tb percentiles during 24 h) (Figure S6d). The phenotype of increased Tb variability was stable through 23 weeks after tamoxifen treatment, the last time studied.

We next explored the response of POA^BRS3^::TeNT mice to various stimuli that increase Tb. The fever response to lipopolysaccharide (LPS) was intact, as was the Tb increase in response to cage switch (Figure 7c, Figure S6f), but not the increase in light-phase Tb produced by a BRS3 agonist (Figure S6e). Thus, POA^BRS3^ neurons are not necessary for the Tb effects of LPS (unlike EP3R/Vglut2-expressing neurons (Machado et al., 2020)) or cage switch. The increased Tb variability of the silenced mice meant these experiments are underpowered to detect if the rise in Tb was augmented.

To explore the response to a Tb-lowering condition, the mice were fasted for 24 h. Both control and silenced mice entered torpor, with the POA^BRS3^::TeNT mice showing an exaggerated hypothermic response (Figure 7d).

We next studied the effect of ambient temperature (Ta). POA^BRS3^::TeNT mice at 22 °C were brought to 30 °C for three days and then returned to 22 °C. There was no major phenotype during the first two stages, but upon return to 22 °C there was a remarkable reduction in Tb (2.8 °C below controls) and energy expenditure (31 % below controls) (Figure 7e,f). We next explored the acclimation in more detail. After acclimation for > 5 days to either 22 °C or 30 °C, mice were acutely exposed (3 h, light phase) to various Ta (Figure 7g). After 22 °C acclimation, the POA^BRS3^::TeNT mice responded to 10-22 °C the same as controls, while at 30 or 34 °C they had a slightly higher Tb. In contrast, when acclimated to 30 °C, the POA^BRS3^::TeNT mice at 16 or 22 °C were much cooler than controls, while still warmer at 34 °C.

Since Tb acclimation to Ta extremes is poorly understood in mice and the POA^BRS3^::TeNT mice provide a robust phenotype, we studied acclimation further. Whether acclimated for 3 or 10 days to 30 °C, the Tb response to 22 °C was similar, taking about 5 days to completely return to baseline (Figure S6g). Thus, these data suggest that acclimation from 22 °C→30 °C occurs by ∼3 days and acclimation from 30 °C→22 °C takes ∼3-5 days.

Taken together, the phenotype of the POA^BRS3^::TeNT mice can be summarized as showing increased Tb variability and exaggerated Tb changes, both increases and decreases, in response to multiple challenges. It appears there is eventually appropriate setting of the target Tb or “set point” in response to various interventions, but poor feedback control of keeping the Tb at the set point.

## Discussion

### Role of POA^BRS3^ neurons in Tb and cardiovascular regulation

We demonstrate that activating POA^BRS3^ neurons increased Tb, HR, and MAP, which is the opposite effect of activating other POA neuron populations, such as those identified by expression of Vglut2/PACAP/leptin receptor (Abbott and Saper, 2017; Hrvatin et al., 2020; Moffitt et al., 2018; Tan et al., 2016; Yu et al., 2016), QRFP (Takahashi et al., 2020), Esr1 (Zhang et al., 2020), Vgat (Zhao et al., 2017), BDNF (Tan et al., 2016), galanin (Kroeger et al., 2018), TRPM2 (Song et al., 2016), NOS1 (Harding et al., 2018), and/or PGDS2 (Wang et al., 2019). There is a POA population expressing Vglut2 and EP3R that mediates LPS-induced fever (Machado et al., 2020). However, PGE2 binding to EP3R inhibits the MnPO^EP3R/Vglut2^ neurons, increasing Tb upon inhibition, which is the reverse polarity of POA^BRS3^ neurons. Thus, the POA^BRS3^ neurons are a novel population that increases Tb when activated.

Activated POA^BRS3^ neurons increase Tb via at least three output pathways (DMH, PVH, and PAG) (Figure S7). Previously, the importance of POA→DMH and POA→RPa pathways had been recognized (Morrison and Nakamura, 2019; Tan and Knight, 2018). Here we add the POA^BRS3^→PVH and POA^BRS3^→PAG pathways to the Tb regulation landscape. The RPa receives input from the DMH and drives preganglionic sympathetic neurons that activate BAT and the cardiovascular system (Cao et al., 2004; Cao and Morrison, 2003), e.g. in response to cold (Nakamura and Morrison, 2007), but POA^BRS3^→RPa activation does not increase Tb. Earlier reports suggest POA→RPa populations are glutamatergic and can express EP3R (Nakamura et al., 2009, Neuroscience; Machado et al., 2020) or QRFP (Takahashi et al., 2020), but activation of the latter did not change Tb. The PVH might increase Tb by direct projections to preganglionic sympathetic neurons (Sutton et al., 2014), however, see (Madden and Morrison, 2009). The PAG output pathway has not been defined. It is also not known if these nuclei contribute equally to other thermoregulatory physiology, such as blood flow redistribution to regulate heat conservation/loss.

The HR response to stimulation of POA^BRS3^→DMH and POA^BRS3^→PVH neurons is mediated by the sympathetic nervous system. Propranolol at 5 mg/kg partially and at 50 mg/kg more completely inhibits sympathetic innervation of the heart (Prando et al., 2018). With a small HR increase persisting during optogenetic stimulation of POA^BRS3^ neurons with 50 mg/kg propranolol, one cannot rule out a small parasympathetic contribution, but the predominant route is clearly sympathetic. As part of the cold-defense response in small mammals such as rats and mice, the HR increases, increasing cardiac output to provide fuel and oxygen to BAT and to distribute heat from BAT to the body, and possibly to directly generate heat from cardiac work. In contrast, mild cold exposure in humans slightly reduces HR, likely via a reflex bradycardia compensating for increased blood pressure due to vasoconstriction (Brychta et al., 2019; Hess et al., 2009). While the POA^BRS3^→RPa activation did not alter Tb, we did not test if this pathway regulates the cardiovascular system.

### Classes of POA^BRS3^ neurons—unraveling the complexity

POA^BRS3^ neurons are not a single population, as analysis of the Moffit et al dataset identifies a minimum of one glutamatergic and five GABAergic POA^BRS3^ populations. Since Vglut2-expressing neurons mediate the metabolic actions of BRS3 (Xiao et al., 2017), we hypothesize that the POA^BRS3^ neurons that increase Tb and HR use glutamatergic transmission. This would be in line with findings showing a glutamatergic POA→DMH pathway involved in cold defense (da Conceicao et al., 2020). Intersectional strategies to selectively manipulate discrete subsets of BRS3 populations (POA^BRS3/Vglut2^ vs POA^BRS3/Vgat2^ and also those expressing other markers of thermogenic neurons--BDNF, PACAP, and/or NOS1) can address this question.

The POA is an anatomically complex region. We optogenetically activated the anteroventral POA that includes the ventral MnPO, VMPO, and VLPO while the chemogenetic experiments additionally included the dorsal MnPO and MPA. These anatomic differences may account for the differences between the optogenetic (increasing Tb) vs chemogenetic (biphasic effect) activation. Further studies are needed to examine anatomical subgroupings of POA^BRS3^ neurons.

Activating POA^BRS3^→PVH, POA^BRS3^→DMH, or POA^BRS3^→PAG neurons each increased Tb. This could be via three distinct neuronal populations projecting to one area each or via one population of neurons having collaterals to three areas. The POA^BRS3^→PVT and POA^BRS3^→RPa populations are distinct, as their activation did not affect Tb. Thus, consideration of projections may also increase the number of POA^BRS3^ classes, and we have studied the function of only five of the output pathways. Investigation is needed to precisely define POA^BRS3^ neuron classes by mRNA expression pattern, neurotransmitter type, cell body location, output field, and physiologic effect.

### Functions of POA^BRS3^ neurons

The POA^BRS3^ neurons increase Tb and HR to contribute to cold defense. The effector pathway for POA neurons to activate BAT has been summarized as a POA→DMH inhibitory pathway, which is inhibited to warm up the animal (Morrison and Nakamura, 2019), although a POA→DMH excitatory pathway also contributes (da Conceicao et al., 2020). One example illustrating the greater underlying complexity is a glutamatergic parathyroid hormone 2 receptor-expressing POA→DMH neuron population that can activate BAT (Dimitrov et al., 2011). The POA^BRS3^ neurons increase Tb by projections to the DMH, PVH, and PAG, adding the PVH and PAG as direct POA thermogenic pathways beyond the DMH and RPa. A prior report proposed that the DMH mediates all autonomic responses evoked from the POA (Hunt et al., 2010). That observation is consistent with the result of optogenetic stimulation of POA^All^ neurons, but is superseded by the more nuanced information from studying neuronal subpopulations such as POA^BRS3^.

The PVH is considered an essential contributor to cold defense, regulating BAT sympathetic activity through projections to the spinal cord intermediolateral column (Amir, 1990; Cano et al., 2003; Saper et al., 1976), but see also (Madden and Morrison, 2009). The PVH also originates sympathetic signaling to other organs, including the heart (Nunn et al., 2011). The POA→PVH pathway has been implicated in autonomic cardiovascular and renal regulation (Frazier et al., 2020; Llewellyn et al., 2012; Marciante et al., 2020; McKinley et al., 2015; Sawchenko and Swanson, 1983; Stocker and Toney, 2005), but less in Tb regulation. We now show that POA^BRS3^→PVH neurons regulate Tb, HR, and MAP through the sympathetic nervous system.

MnPO→PAG neurons are activated by various challenges (Uschakov et al., 2009; Yoshida et al., 2005): neurons in the PAG can activate BAT (Cano et al., 2003; Chen et al., 2002) and stimulation of hypothalamic projections to PAG increased Tb (de Git et al., 2018). We now report that selectively stimulating POA^BRS3^→PAG projections increased Tb. More experiments will further establish the function of this pathway.

Together, we establish BRS3 as a marker for the thermogenic POA→DMH pathway and add two new pathways, POA^BRS3^→PVH and POA^BRS3^→PAG, that can increase Tb when activated. All possibly contribute to the cold defense response.

### POA^BRS3^ neurons fine-tune regulation of Tb

POA^BRS3^ neurons are not merely upstream drivers of BAT activation. They also provide the interesting function of fine-tuning Tb regulation. When POA^BRS3^ neurons were constitutively silenced, Tb was more variable with a tendency to bidirectionally overshoot the Tb of the controls. Although non-significant in some cases, mice with silenced POA^BRS3^ neurons generally had a lower Tb during times with lower Tb (fasting-induced hypothermia, cold Ta, light phase) and a higher Tb during times with higher Tb (hot Ta, dark phase, handling, LPS treatment, BRS3 agonist treatment).

Taking advantage of the robust signal in POA^BRS3^ silenced mice, we studied the timing of central thermal acclimation. In the silenced mice, Tb acclimation occurred by ∼3 days after shifting from 22 °C→30 °C, while acclimation from 30 °C→22 °C took ∼3-5 days. To our knowledge the timing of central thermal adaptation in mice has not been reported previously. Longer times, 1-2 weeks, are required for full remodeling of peripheral BAT in response to cold exposure (Cinti, 2009).

It is notable that in each situation studied, the silenced mice made the physiologically appropriate underlying response to the intervention, albeit accompanied by overshoot and increased variability. The mice did not overcome this mis-regulation - it was stable over the 6 months they were studied. Since the Tb variability of *Brs3^-/y^* mice is not increased (Lateef et al., 2014), this effect is a property of POA^BRS3^ neurons and not due to a deficit in BRS3 signaling *per se*.

In other examples of Tb overshoot, the overshoot is not bidirectional. Mice with ablated MnPO^Vgat^ or MnPO^Vglut2^ neurons have the same or higher Tb in both hot and cold environments (Machado et al., 2020). Mice lacking adipose tissue (Gavrilova et al., 1999), overexpressing FGF21 (Inagaki et al., 2007), or with ablated orexin neurons (Futatsuki et al., 2018) have lower Tb upon challenge with cold or fasting, while none appears to overshoot an increased Tb situation. Thus, one possibility is that the overshoots in POA^BRS3^ silenced mice are due to silencing of two opposing POA^BRS3^ neuronal populations. This is supported by the faster return to baseline in our chemogenetic activation data. Furthermore, some POA BRS3 populations also express PTGDS, BDNF, PACAP, ER1a, NOS1 and galanin, all markers for neurons driving hypothermia when activated and/or involved in the heat or torpor response.

A different possible mechanistic explanation for the overshoot is impaired sensation and evaluation of the internal and/or external environment. Multiple ion channels contribute to sensing internal and external temperature. *Trpv1^-/-^;Trpm8^-/-^;Trpa1^-/-^* mice, which lack three of these channels, have remarkably normal thermal biology and Tb variability (Skop et al., 2020a). In contrast, mice with neonatal ablation of sensory neurons with resiniferatoxin do show increased Tb variability and their Tb is even more poikilothermic (lower in a cold Ta and higher in a warm Ta) than the POA^BRS3^ silenced mice (Skop et al., 2020a). It is possible that POA^BRS3^ neurons integrate sensory information for Tb regulation. Along these lines, another possibility for the increased Tb variability is that the thermal sensory information from the LPB is not properly integrated in the POA integrative thermoregulatory circuitry when the POA^BRS3^ neurons are permanently silenced. This could make the system more labile.

BRS3 is a marker for multiple neuronal populations in the POA that are involved in increasing Tb, HR, and MAP via sympathetic nerves. POA^BRS3^ neurons contribute to cold defense and fine-tuning of Tb regulation. Further identification of POA^BRS3^ neuron subclasses, and their connectivity and function, will advance our understanding of homeothermy, a defining feature of mammalian biology.

### Limitations of Study

One limitation is the existence of multiple POA^BRS3^ neuronal populations. Thus, further studies are needed to identify molecular markers allowing selective study of POA^BRS3^ subpopulations, with elucidation of each neuronal subpopulation’s neurotransmitters, functions, inputs, and projections. For instance, an unanswered question is if the thermogenic POA^BRS3^ neurons are glutamatergic, GABAergic or both. Also, the identities of the downstream neurons in PVH and DMH and their pathways are need for a fuller picture of the circuits. Finally, we have not performed in vivo imaging to correlate neuronal activation with specific thermal biology.

## Acknowledgments

We thank Alice Franks for superb technical assistance, Yuning Huang for E-Mitter implantation, Naili Liu for indirect calorimetry, and Audrey Noguchi and Danielle Springer of the NHLBI Murine Phenotyping Core for the cardiovascular telemetry implantation surgeries. This research was supported by the Intramural Research Program of the National Institutes of Health, National Institute of Diabetes and Digestive and Kidney Diseases (ZIA DK075062; ZIA DK075063, ZIA DK075064, ZIA DK070002, ZIA DK075087, ZIA DK075088).

## Author contributions

RAP and MLR conceived and designed the study with input from MJK, OG, and CX. RAP performed and analyzed the experiments. RAP, ASM, CKH and AS performed optogenetic experiments, immunohistochemistry and counted cells. RAP, CKH and AS performed tracing experiments. OG performed indirect calorimetry experiments. VS and RAP performed IR experiments. CL and RAP performed electrophysiology experiments. CKH analyzed expression profiling data. ASM, RAP, and HSP performed chronic silencing experiments. RAP wrote the manuscript with input from MLR and all other authors.

## Methods

### Animals

Animal studies were approved by the NIDDK/NIH Animal Care and Use Committee (protocol K016-DEOB-20). Mice were singly housed after surgery at 21-22 °C with lights on 6 am–6 pm. Chow (7022, Envigo, Indianapolis, IN) and water were available ad libitum, including during drug treatments, indirect calorimetry, and optogenetic experiments, with exception of infrared thermography and food intake experiments as indicated. Mice used were: Brs3-2A-CreER^T2^ (BRS3-Cre; Jax# 032614) (Pinol et al., 2018), B6.Cg-Gt.(Rosa)26Sortm6(CAG-ZsGreen1)Hze/J, with Cre-dependent ZsGreen (Ai6; Jax# 007906) (Madisen et al., 2010), B6.Cg-Gt(ROSA)26Sortm14(CAG-tdTomato)Hze/J, with Cre-dependent tdTomato (Ai14; Jax# 007914) (Madisen et al., 2010), Gad2-2a-NLS-mCherry (Gad2-mCherry; Jax# 023140)(Peron et al., 2015), and C57BL/6J (Jax# 000664). *Brs3* is on the X chromosome and male mice were used, unless otherwise noted. For all behavioral and physiological studies, male mice were between 8 and 45 weeks of age. Mice for ChR2-assisted circuit mapping (CRACM) studies (male) were between 5 and 13 weeks of age. Mice were on a mixed C57BL/6J and 129SvEv background. Mice from multiple litters were used for all studies.

*Tamoxifen.* Cre-mediated recombination was achieved by treatment with tamoxifen (75-110 mg/kg i.p. in corn oil; Sigma Aldrich) daily for 5 consecutive days, with mice studied ≥4 weeks after the first dose.

### Stereotaxic injections

*General surgical procedures*. Mice were anesthetized with 0.5-1.5% isoflurane (1 L/min oxygen) or a ketamine/xylazine mix (80/10 mg/kg, i.p.), placed in a stereotaxic instrument (Digital Just for Mouse Stereotaxic Instrument, Stoelting), and ophthalmic ointment (Puralube, Dechra) was applied. Brain injections were done with pulled-glass pipettes (pulled 20–40 µm tip diameter; 0.275 mm ID, 1 mm OD, Wilmad Lab Glass) at a visually controlled rate of ∼50 nl per min with an air MAP system regulator (Grass Technologies, Model S48 Stimulator). The pipette was kept in place for 5 min after injection. Post-surgery, mice received subcutaneous saline injections to prevent dehydration and analgesic (buprenorphine, 0.1 mg/kg, s.c.).

*Viral vectors*. We used the following viruses: AAV1-ef1a-DIO-ChR2-EYFP (gift from K. Deisseroth; Addgene viral prep 20298-AAV1); AAV9-Ef1a-DO-hChR2(H134R)-mCherry (gift from B. Sabatini; Addgene plasmid 37082(Saunders et al., 2012); Vigene Biosciences, Inc.); AAV8-Ef1a-DIO-synaptophysin-mCherry (Virovek, Inc.); AAV-DJ-CMV-DIO-eGFP-2A-TeNT (Stanford Viral Core GVVCAAV-71; Similar to (Campos et al., 2018)); AAV5-EF1a-DIO-EYFP-WPRE-pA (UNC Vector Core); pAAV8-hSyn-DIO-hM4D(Gi)-mCherry (gift from B. Roth; Addgene viral prep # 44362-AAV8), pAAV8-hSyn-DIO-hM3D(Gq)-mCherry (gift from B. Roth; Addgene viral prep 44361-AAV8), AAV8-Ef1a-DIO-mCherry (gift from B. Roth; Addgene viral prep # 50459-AAV8)(Krashes et al., 2011); AAV8-Ef1a-FLEX-TVA-mCherry (UNC vector core); EnvA-G-Deleted-Rabies-Egfp (Gene Transfer, Targeting and Therapeutics Core at Salk Institute).

### Immunohistochemistry

*Tissue Preparation.* Mice were anesthetized (chloral hydrate, 500 mg/kg, i.p.), perfused transcardially with 0.9% saline followed by 10% neutral buffered formalin, and the brain was removed. Brains were post-fixed (10% formalin, room temperature, overnight) and incubated in 30% sucrose in 0.1M PBS (4⁰C, overnight or until use), sectioned coronally into three equal series (50 μm sections) on a sliding microtome (SM2010 R, Leica) and collected in 0.1M PBS. Sections were washed 3×10 minutes in 0.1M PBS and the following steps were performed with shaking at room temperature and in 0.1M PBS solutions, unless noted otherwise: *mCherry.* Sections were incubated for 1 hour in 0.3% Triton X-100, 3% normal goat serum (NGS) and then incubated overnight with 1:2000 rabbit-anti-DsRed (632496, Clontech) (Geerling et al., 2017) in 0.3% Triton X-100, 3% NGS. Sections were washed, incubated (2 hours) with secondary antibody (1:500 Alexa-555 goat-anti-rabbit; A-21428, Thermo Fisher) in 0.3% Triton X-100 and 2% NGS.

*GFP.* Sections were processed as for mCherry, using 1:1000 chicken-anti-GFP (13970, Abcam) and 1:500 Alexa-488 goat-anti-chicken (A-11039, Thermo Fisher) as the primary and secondary antibodies.

After incubation with secondary antibody, sections were washed, mounted and cover slipped with Prolong mounting medium containing DAPI (Thermo Fisher).

### Optogenetics

*Virus injections and fiber implantation*. BRS3-Cre or BRS3-Cre;Ai14 mice were injected with 50 nl AAV1-Ef1a-DIO-ChR2-EYFP, AAV9-Ef1a-DIO-hChR2(H134R)-mCherry, AAV9-Ef1a-DO-hChR2(H134R)-mCherry or, as controls, AAV8-Ef1a-DIO-mCherry in the anteroventral POA (AP: 0.35; ML: 0.3; DV: −5.25, mm from bregma). Following virus injection, optical fibers (200 μm diameter core; NA 0.22; Nufern), glued to ceramic zirconia ferrules (230 μm bore; 1.25 OD diameter; Precision Fiber Products), were implanted unilaterally over the anteroventral POA (AP: 0.35; ML: 0.3; DV: −4.5, mm from bregma), PVH (AP: −0.9; ML: 0.25; DV: −4.5, mm from bregma), DMH (AP: −1.85; ML: 0.3; DV: −4.5, mm from bregma), PVT (AP: −1.85; ML: 0.3; DV: −2.2, mm from bregma), RPa (AP: −6.1; ML: 0; DV; −5.50, mm from bregma) or PAG (AP: −4.5; ML: 0.5; DV: −2.0, mm from bregma). Fibers were fixed to the skull using C&B Metabond Quick Cement and dental acrylic.

*Experimental procedures*. Mice were allowed to adapt to the fiber patch cord for at least two days prior to experiments and typically not handled on the day of the experiment. Fiber optic cables (200 µm diameter, NA 0.22, 1 m long, Doric Lenses; or, 0.5m long, ThorLabs) were connected to the implanted fiber optic cannulas with zirconia sleeves (Precision Fiber Products) and coupled to lasers via a fiber optic rotary joint (Doric Lenses). We adjusted the light power of the laser (473 nm; Laserglow or Opto Engine) such that the light power (measured with a fiber optic power meter; PM20A; ThorLabs) at the end of the fiber optic cable was ∼10 mW. Using an online light transmission calculator for brain tissue (https://web.stanford.edu/group/dlab/cgi-bin/graph/chart.php), we estimated the light power at tip of implanted fiber between 3 and 6 mW/mm^2^. This is an upper limit due to possible light loss between the fiber optic cable and the implanted optic fiber. Light pulses were controlled by a programmable waveform generator (Arduino). Pulses were 10 ms delivered at 20 Hz and stimulation was on for 1 s, followed by 3 s off (POA and IR study) or 1 s on/1 s off (POA^BRS3^ projections) or continuous (short stimulations in POA). After completion of experiments, fiber placement and ChR2 expression were assessed. Animals without ChR2 expression or incorrect placement of optic fibers were excluded from analysis.

*Analysis:* Within each mouse, the multiple optogenetic stimulation epochs (5-10) were normalized to the baseline immediately before laser on and averaged. Because of the body’s heat capacity, Tb changes are gradual (unlike HR and MAP), so we used 10 to 19 min from laser on as the Tb response period, but 0 to 9 min for HR, MAP, and physical activity. Response period is indicated as On. Off is −10 to −1 min before laser On.

### Chemogenetics experiments

*POA virus injection*. BRS3-Cre or BRS3-Cre;Ai6 mice received bilateral injections of 50 nl AAV8-hSyn-DIO-hM3Dq-mCherry or AAV8-hSyn-DIO-hM4Di-mCherry in the anterior POA (AP: +0.35; ML: +/- 0.30; DV: −5.30, mm from bregma). Expression was verified after IHC for mCherry. All included POA^BRS3^::hM3Dq mice had bilateral expression in MnPO, VMPO, and MPA.

POA^BRS3^::hM4Di mice with uni- and bilateral expression in the POA (anterior and intermediate) were used. Mice without hM4Di-mCherry expression were excluded.

*BNST virus injection*. BRS3-Cre or BRS3-Cre;Ai6 mice received bilateral injections of 50 nl AAV8-hSyn-DIO-hM3Dq-mCherry or AAV8-hSyn-DIO-mCherry in the BNST (AP: −0.22; ML: +/- 0.85; DV: −3.75, mm from bregma). Mice with uni- and bilateral hits were used. Mice without hM3Dq-mCherry expression were excluded.

*Tb telemetry.* Tb experiments were performed described as below and in legends. The return to baseline Tb was measured as the minute the Tb is within 0.05 °C of the mean baseline (−150 to − 30 min). One mouse consistently did not return to baseline and was excluded from analysis.

*Food intake inhibition.* Mice were fasted five hours before the onset of their dark cycle. Mice were dosed with CNO or vehicle 15 minutes prior to lights out. Food intake was measured two hours following dosing.

*Food intake stimulation*. Ad-lib fed mice were dosed with CNO or vehicle four hours into the light cycle and food was removed. After 15 minutes food was returned and food intake was measured over the next two hours.

### Body temperature telemetry

Animals were anesthetized as above and E-Mitters (Starr Life Sciences) were implanted intraperitonially (Lute et al., 2014). At least two days before experiments, mice were housed in temperature-controlled chamber in their home cages on ER4000 energizer/receivers (Starr Life Sciences). Tb and activity were continuously measured by telemetry with 1-min means collected with VitalView software (Starr Life Sciences). Experiments were performed at 22 °C, unless indicated otherwise. Physical activity is measured in arbitrary units.

### Blood pressure and heart rate telemetry

In a subset of mice with a Tb response to optogenetic stimulation, we replaced the E-Mitter with an intra-arterial pressure telemetry probe (model HD- X10 or HD-X11; Data Sciences International, St Paul, MN) and continuous ambulatory blood pressure, heart rate, physical activity and subcutaneous (HD-X10) or intraperitoneal (HD-X11) Tb were measured (Kim et al., 2008). Data were sampled at 1000 Hz, processed using a PhysioTel RPC-1 receiver, and collected with Ponemah v6.30 (Data Sciences International). Unless noted, 1 min averages were used for analysis. Physical activity is measured in arbitrary counts.

### Drugs

Lipopolysaccharide from Salmonella enterica serotype typhimurium (LPS; Sigma) was dissolved in sterile saline and administered at 100 µg/kg i.p. MK-5046 (Sebhat et al., 2011) (MedChemExpress, Monmouth Junction, NJ) was dissolved in vehicle of 10% Tween 80/0.25% methylcellulose in saline and administered at 10 mg/kg i.p. Propranolol is a β-adrenergic antagonist that has ∼100-fold selectivity for β1 and β2 over β3 adrenergic receptors (Baker, 2005; Cernecka et al., 2014; Popp et al., 2004). Propranolol (Sigma) was dosed i.p. at 0-100 mg/kg (vehicle was 10% DMSO for 30 - 100 mg/kg; water for lower doses) for the Tb dose response. For the cardiovascular response propranolol (vehicle, 10% DMSO) was injected at 0, 5, or 50 mg/kg, i.p., at 5 µl/g.

### Anterograde and collateral tracing

*Anterograde tracing.* BRS3-Cre or BRS3-Cre;Ai6 mice were injected with 25 nl of AAV8-Ef1a-DIO-synaptophysin-mCherry in the anteroventral POA (AP: 0.35; ML: 0.3; DV: −5.25, mm from bregma). Mice were euthanized 5-8 weeks after start of tamoxifen treatment and brains processed for immunohistochemistry (see above). All 5 mice with expression limited to the anterior POA had consistent axonal projections to described areas. *POA^BRS3^→DMH collateral tracing* AAV-DIO-TVA-mCherry was unilaterally injected in the POA of BRS3-Cre mice (50 nl). >4 weeks after tamoxifen treatment retrograde G-deleted-Rabies-GFP (100 nl) was (ipsilateral as the first viral injection) injected in the DMH. Mice were perfused 12d after last injection and brains processed for IHC for GFP and mCherry.

### Quantitative thermal imaging in freely moving mice

Shaved skin temperatures in the interscapular and dorsal lumbar regions were used as measures of BAT and core body temperature, respectively (Gachkar et al., 2017; Pinol et al., 2018; Vianna and Carrive, 2012) (Skop et al., 2020b). In brief, one day prior to study, the interscapular region and a 2 x 2 cm midline area 2 cm above the tail were shaved under isoflurane anesthesia. After housing overnight in their home cage with the optical fiber attached, mice were placed in a 20 x 20 cm enclosure with bedding, 170 cm below the infrared (IR) camera (T650sc, FLIR Systems, Wilsonville, OR), and acclimated for 3 h. Mice then underwent 3 cycles of 20 min baseline, 20 min laser on, and 40 min post-stimulation, with IR images collected continuously using ResearchIR software (FLIR Systems, Wilsonville, OR; 7.5 frames per second; IR emissivity set to 0.97 (Pinol et al., 2018)). Maximum interscapular and lumbar temperatures were determined by a blinded observer every 2 minutes using FLIR Tools® (FLIR Systems, Wilsonville, OR). Tail temperature was measured 1 cm from the body at baseline (0 to 20 minutes before stimulation onset) and during stimulation (6 to 10 minutes after stimulation onset). A limit of this assay is that the mice need to be awake and moving for the camera to capture the tail. The ambient temperature was 25-26 °C. Infrared data were exported as a video file, from which physical activity was determined by EthoVision XT software (Noldus, Leesburg, VA).

### Neurochemical identity

We bred triple transgenic mice (BRS3-Cre;Ai6;Gad2-mCherry) in which Gad2 neurons express mCherry and BRS3 neurons express ZsGreen. We performed immunohistochemistry for mCherry and counted the colocalization of BRS3-positive neurons and Gad2-positive neurons in every third coronal brain section. Data are presented as mean with individual mice.

### Electrophysiology

BRS3-Cre mice (5-10 weeks) received bilateral 50 nl injections of AAV1-Ef1a-DIO-ChR2-EYFP in the anteroventral POA (AP: 0.5; ML: +/- 0.3; DV: −5.25, mm from bregma). Four to 6 weeks later, brain slices were obtained and stored at 30 °C in a heated, oxygenated chamber containing aCSF (in mmol/l) 124 NaCl, 4.4 KCl, 2 CaCl_2_, 1.2 MgSO_4_, 1 NaH_2_PO_4_, 10.0 glucose, and 26.0 sodium bicarbonate before being transferred to a submerged recording chamber maintained at 30 °C (Warner Instruments, Hamden, CT). Recording electrodes (3-5 MΩ) were pulled with a Flaming-Brown Micropipette Puller (Sutter Instruments, Novato, CA) using thin-walled borosilicate glass capillaries.

Light evoked excitatory and inhibitory postsynaptic currents (EPSCs and IPSCs, respectively) were measured in voltage-clamp mode using electrodes filled with an intracellular recording solution containing (in mM): 135 Cs-methanesulfonate, 10 KCl, 10 HEPES, 1 MgCl_2_, 0.2 EGTA, 4 Mg-ATP, 0.3 GTP, 20 phosphocreatine, 2 QX314. Neurons were held at −55 mV to isolate glutamatergic synaptic transmission and record EPSCs, or +10 mV to isolate GABAergic synaptic transmission and record spontaneous IPSCs within individual neurons. Tetrodotoxin (TTX, 500 nM) and 4-aminopyridine (4-AP, 100 µM) were included in the bath aCSF.

### Tetanus toxin (TeNT) treatment

*AAV injections and expression.* TeNT cleaves synaptobrevin, leaving neurons viable but permanently unable to release neurotransmitters (Sweeney et al., 1995). BRS3-Cre or BRS3-Cre;Ai14 mice received 50 nl bilateral injection of AAV-DJ-CMV-DIO-eGFP-2A-TeNT (control: AAV5-EF1a-DIO-EYFP-WPRE-pA) into the POA (AP: 0.35; ML: 0.3; DV: −5.25, mm from bregma) to chronically silence, but not kill, POA^BRS3^ neurons.

They were also implanted with E-Mitters (see above). After completion of the experiments, brains were processed for immunohistochemistry for GFP and histological analysis. GFP-TeNT-expressing neurons were counted in every third 50 µm coronal section in POA (median preoptic area (MnPO), ventromedial preoptic area (VMPO), and ventrolateral preoptic area (VLPO)) in 4 sections per mouse.

*Fasting.* Mice in their home cages were food-deprived for 24h, starting just before the onset of the dark cycle.

*Responses to acute Ta changes*. Tb and physical activity were recorded in response to acute 3 h Ta changes (last 2 h used for analysis) after >3d adaptation to regular housing Ta (22° C) or a warmer Ta (30° C).

*Drug studies*. Drug or respective vehicles were administered 4-6 h into the light cycle in a crossover design.

*Cage switch.* Mice were switched to a clean cage, which causes a stress response (Balcombe et al., 2004; Rasmussen et al., 2011).

### Indirect calorimetry

An Oxymax/CLAMS (Columbus Instruments) was used to measure Tb, total energy expenditure (TEE), RER (respiratory exchange ratio, O_2_ consumed:CO_2_ produced), and activity by beam break simultaneously in mice implanted with E-Mitters (Lute et al., 2014). Mice were acclimated in the chambers for 2 days before start of the experiment. Experiments were performed at 22 °C and 30 °C, as indicated. Sampling was every 13 minutes, measuring from 12 chambers.

### scRNA analysis

The scRNA seq count matrix for GSE113576 (Moffitt et al., 2018) was analyzed with R (v. 4.0.3) using Seurat (v.3.2.2) (Butler et al., 2018; Stuart et al., 2019). Raw counts were normalized, scaled, and the 2,000 most variable genes were used as input for principal component (PC) analysis. Cells were then clustered (PC: 75, resolution: 1.2) and visualized with t-SNE. Clusters enriched for neuronal markers (Snap25, Eno2, Tubb3, Stmn2) and de-enriched for non-neuronal markers were selected for downstream analysis. Of the 18,401 neuronal cells, we re-clustered the 581 neurons with ≥1 UMI corresponding to Brs3 using the same pipeline as above (PC: 15, resolution: 0.5). Differentially expressed genes (DEGs) and cluster marker genes were identified using the Wilcoxon Rank Sum test. Cluster names were assigned with one significantly enriched marker gene (average log2 fold change ≥0.75 and FDR adjusted p-value ≤1.238E-20). Clusters labeled beginning with ‘i’ represent inhibitory clusters enriched with Vgat, while that labeled with ‘e’ represents an excitatory cluster enriched with Vglut2. The cluster labeled ‘Apoptotic’ is enriched for several mitochondrial genes, suggesting cell death.

### Image capture and processing

*General procedure.* Overview images to verify optic fiber placement and viral expression were captured using an Olympus BX61 motorized microscope with Olympus BX-UCB hardware (VS120 slide scanner) and processed using Olympus OlyVIA software (Olympus). Confocal imaging for images of viral expression and optic fiber placement and quantification of BRS3-positive/GAD2-positive and TeNT-EGFP neurons was performed with an upright Zeiss Axio Observer Z1 microscope with a 10X objective, Zeiss 700 confocal hardware, and Zen software (2012; Zeiss). Images were minimally processed to adjust brightness and contrast.

*Analysis.* Images of every third 50 μm coronal brain section were acquired and neuron counting and colocalization analysis was performed using neuroanatomical landmarks (Franklin and Paxinos, 2007). Neurons were counted manually with the experimenter blinded to the experimental group.

## QUANTIFICATION AND STATISTICAL ANALYSIS

All data are presented as mean ± SEM or mean + SEM, unless otherwise indicated. No statistical methods were used to pre-determine sample size but our sample sizes are similar to those reported in previous publications (Lute et al., 2014). Data distribution was assumed to be normal, but this was not formally tested in experiments other than the permanent inactivation TeNT experiment. The TeNT mice group had higher variance (Figure S6d, bottom panels) and in comparing TeNT and CTRL mice an unequal variance t-test was used. All t-tests were two-sided. All n numbers refer to number of mice used in the experiment; in some experiments, as indicated, data are the average of multiple observations per mouse. Data collection and analysis were not performed blind to the conditions of the experiments, except for IR camera imaging.

We used paired t-tests (2-sided) for within-mouse comparison of the effect of stimulation or unpaired when comparing two groups (e.g. TeNT vs Control). For comparisons between more than 2 groups, we used a one-way ANOVA with multiple comparison testing.

---

**Supplemental Figure 1.**
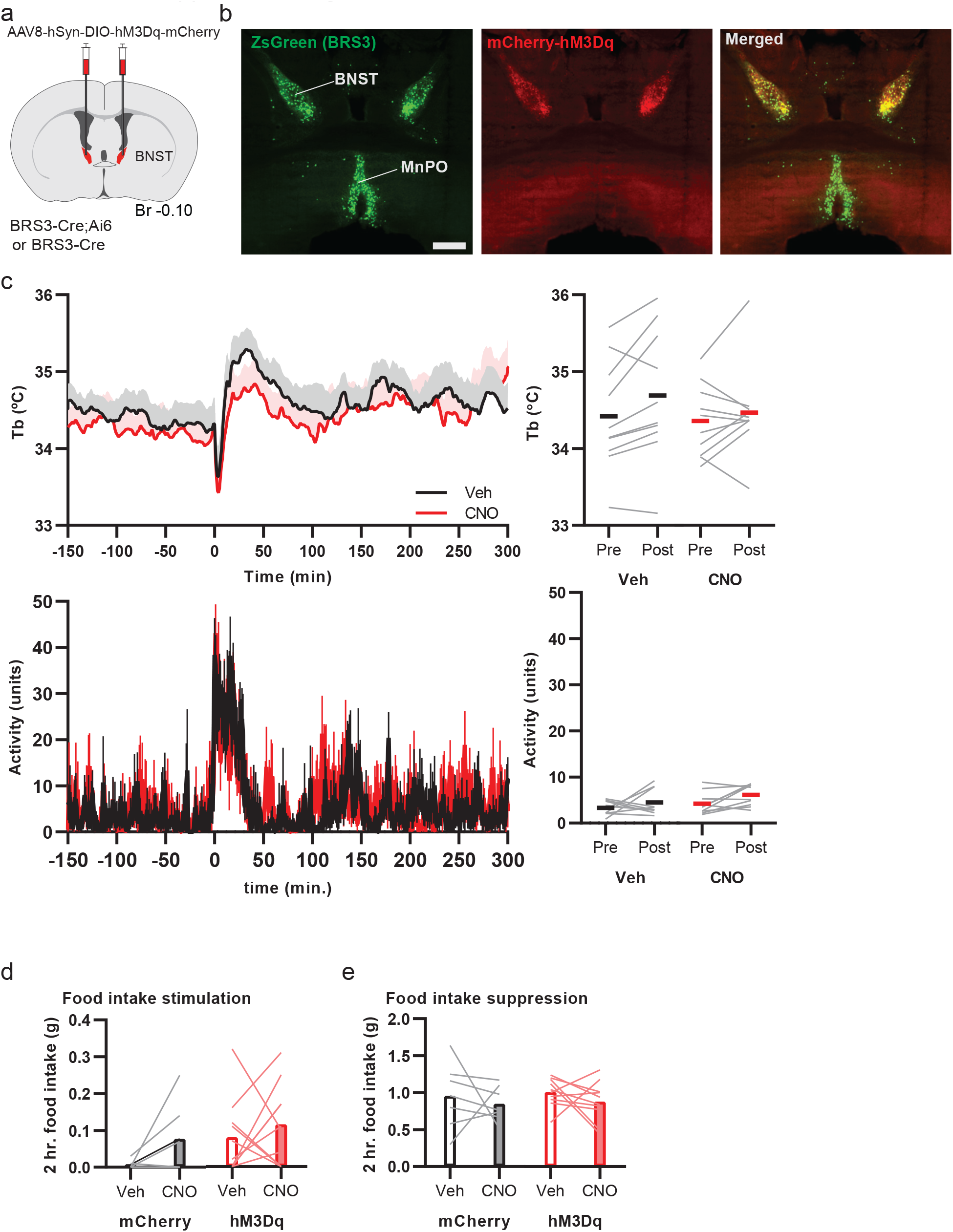
Chemogenetic activation of BNST^BRS3^ neurons does not change Tb, physical activity or food intake. Related to Figure 3. a) Schematic of virus injection. b) hM3Dq expression (red) in BRS3 (green) neurons in the BNST of a BRS3-Cre;Ai14 mouse. Angle of coronal slice was adjusted to include the MnPO and BNST in the same slice. Scale bar is 500 µm c,d) Tb and physical activity response to CNO (1 mg/kg) or vehicle in BNST^BRS3^::hM3Dq mice (n = 10). For quantification, Pre is mean from -150 to −30 and Post from 60 to 180 min. No significant difference in paired two-sided t test on change from baseline, CNO vs vehicle. Data are mean + s.e.m. in top left panel. S.e.m. not graphed for visual clarity in bottom left. Black (vehicle) and red (CNO) bars represent means. d) Effect of CNO (1 mg/kg) on food intake at onset of light cycle in satiated BNST^BRS3^::hM3Dq (n = 10/group) and control BNST^BRS3^::mCherry mice (n = 7/group). e) Effect of CNO (1 mg/kg) on food intake at onset of dark cycle after 5 h fast in BNST^BRS3^::hM3Dq mice (n = 10/group) and control BNST^BRS3^::mCherry mice (n = 7/group). (c-e) Crossover design.

**Supplemental Figure 2.**
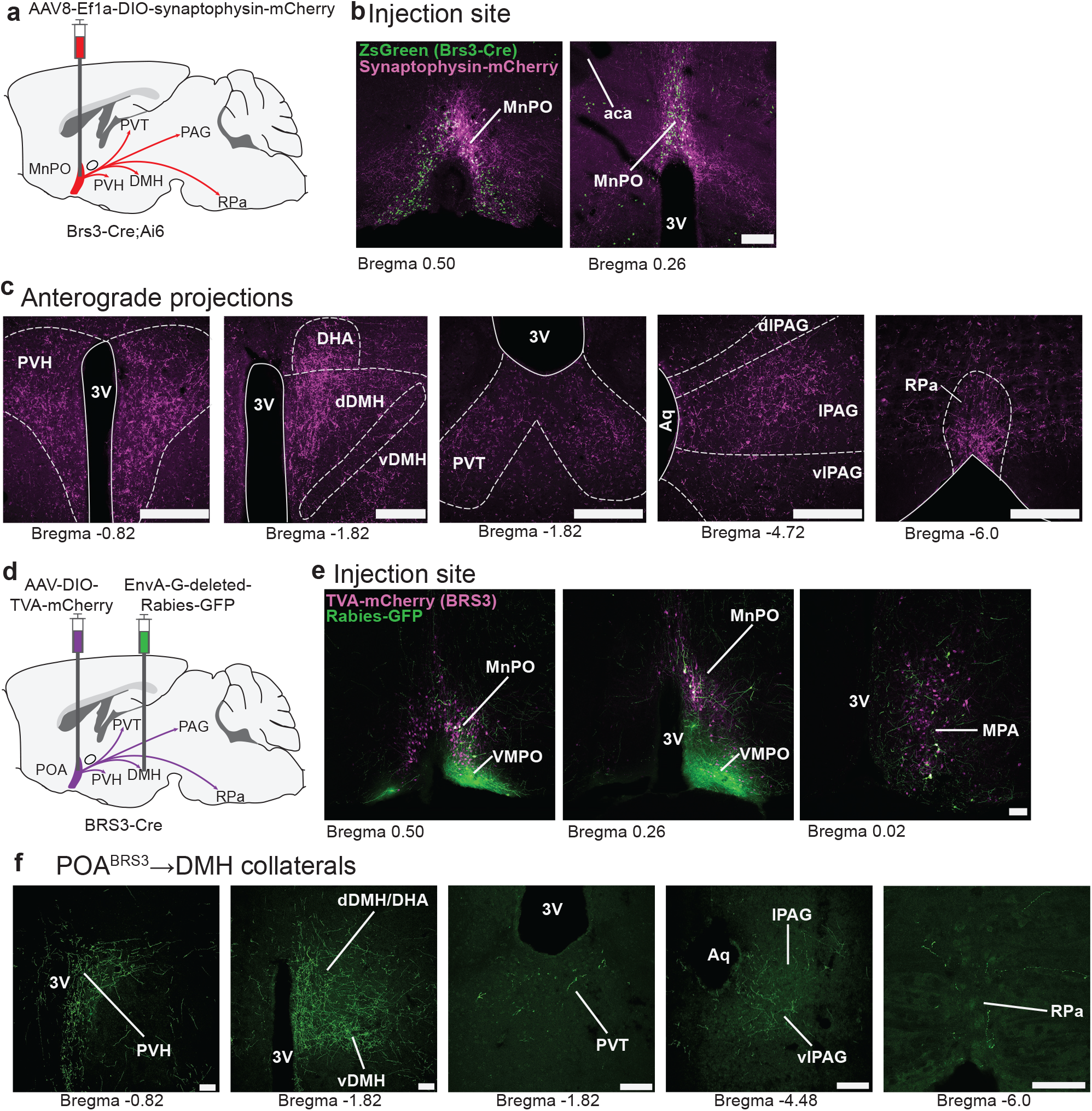
POA^BRS3^ neurons project widely in the brain with collaterals. Related to Figure 4. a) Schematic of virus injection into BRS3-Cre;Ai14 mice for Cre-dependent synaptophysin mCherry expression and the major projections. b) Injection site verification showing that virus was limited to MnPO region of POA. BRS3 neurons are green and viral synaptophysin is magenta. c) Projection targets of MnPO^BRS3^ neurons. b,c) Scale bar is 200 µm. d) schematic of virus injection strategy in BRS3-Cre mice; Rabies-GFP was injected in the DMH >4 weeks after first injections and tamoxifen treatment. e) POA^BRS3^ TVA-mCherry expressing neurons (magenta) and Rabies-GFP-expressing neurons (green), retrogradely infected through axon projections to DMH. f) Projection targets of POA^BRS3^→DMH neurons. e,f) Scale bar is 100 µm 3V – third ventricle; aca – anterior commissure; Aq – aqueduct; dDMH/DHA – dorsal part of the dorsomedial hypothalamus/dorsal hypothalamic area; dlPAG - dorsolateral periaqueductal grey; MnPO – median preoptic area; MPA – medial preoptic area; PVH – paraventricular nucleus of the hypothalamus; PVT – paraventricular nucleus of the thalamus; lPAG - lateral periaqueductal grey; RPa – raphe pallidus; vDMH – ventral part of the dorsomedial hypothalamus; vlPAG – ventrolateral periaqueductal grey; VMPO – ventromedial preoptic area.

**Supplemental Figure 3.**
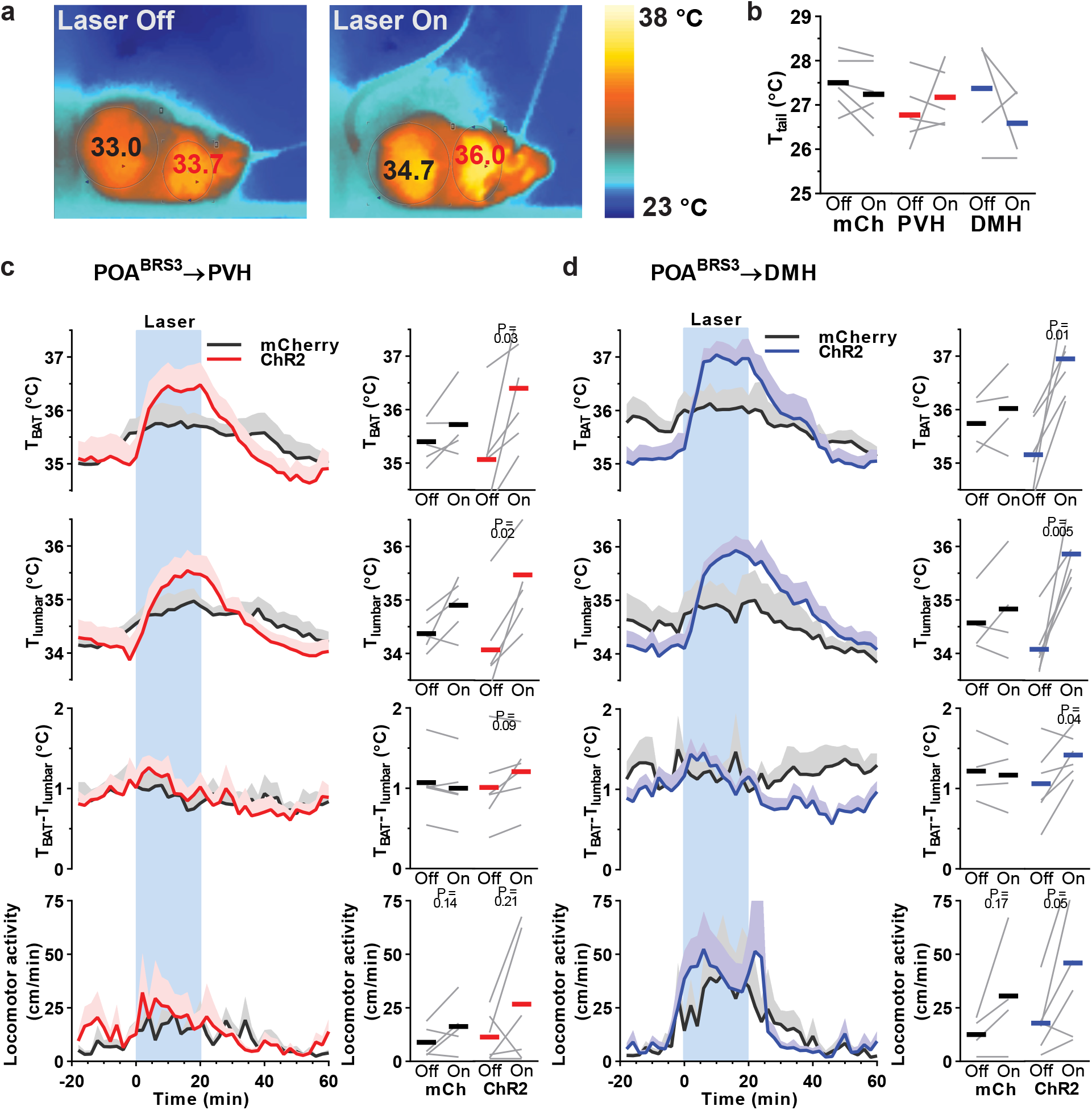
POA^BRS3^→PVH and POA^BRS3^→DMH neurons increase Tb through BAT activation. Related to Figure 4. a) POA^BRS3^→PVH mouse with laser off (left) and on for 6 minutes (right). Infrared camera interscapular (T_BAT_; red) and lumbar (T_lumbar_; black) skin temperature. b) Mean (bars) and individual (gray lines) tail temperature (T_tail_) before and during stimulation. POA^BRS3^→PVH::mCherry and POA^BRS3^→DMH::mCherry mice are combined in the control group. c,d) T_BAT_, T_lumbar_, their difference (T_BAT_-T_lumbar_), and physical activity during optogenetic stimulation (blue interval; 1s on 3s off; 20 Hz; 10 ms pulses) of POA^BRS3^→PVH (c, red, n= 6) and POA^BRS3^→DMH (d, blue, n= 6) projections and respective mCherry controls (black, n = 4-5). Data are average of 3 epochs/mouse, relative to epoch baseline (−20 to −1 min); mean ± s.e.m Quantitation in right panels uses intervals: Off, −10 to −2 min; On, 10 to 18 min for T_BAT_ and T_lumbar_, 0 to 8 min for locomotor activity, and Off, −4 to 0 min; On, 2 to 6 min for T_BAT_-T_lumbar_. Experiments were performed at 25 °C. Bars are means; gray lines, individual animals; P values from paired t test, Off vs On.

**Supplemental Figure 4.**
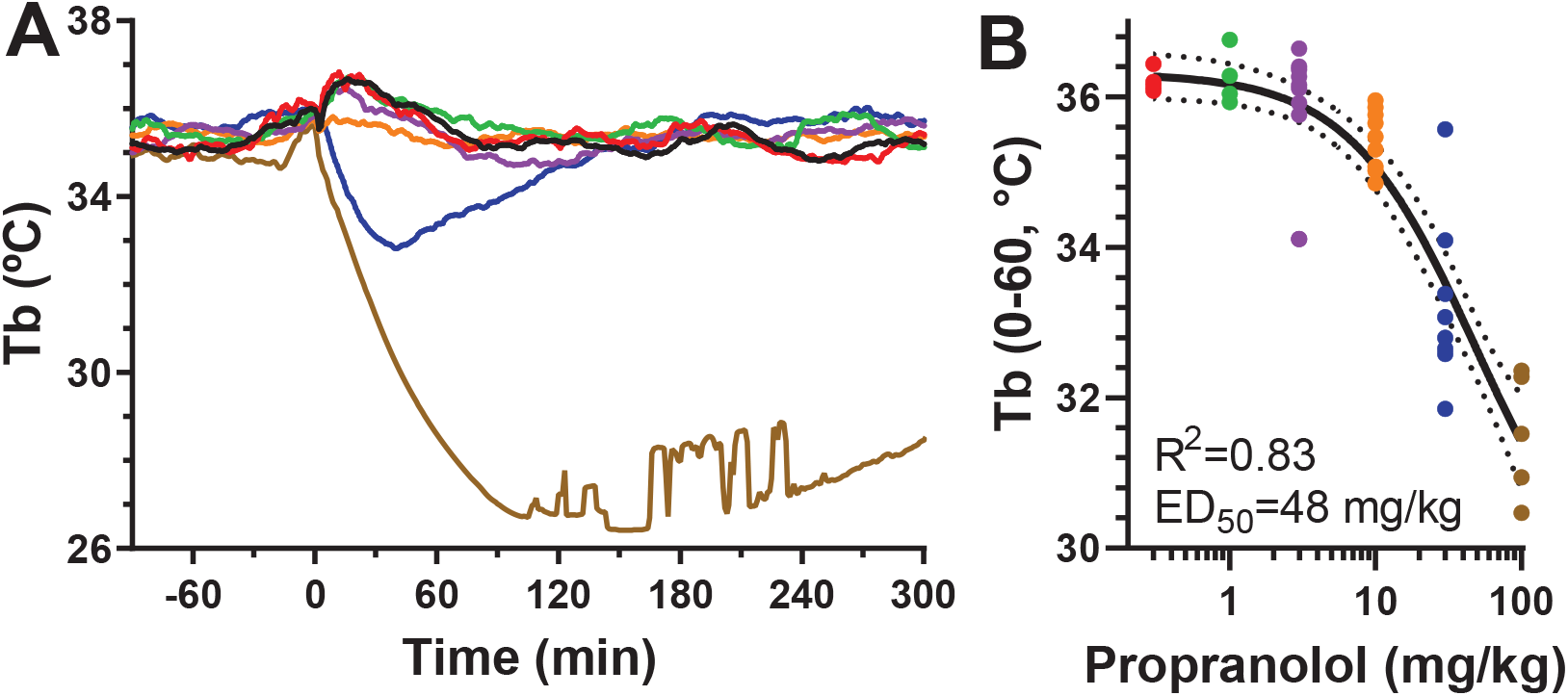
Effect of propranolol on body temperature. Related to Figure 4. a) Wild-type mice were treated with the indicated dose of propranolol (color key in b) or vehicle (10% DMSO for 30 and 100 mg/kg; water for other doses) at time 0. b) Tb (mean of 0-60 min after dosing) data were non-linearly fit using Prism to Tb = bottom+(top-bottom)/(1+(dose/ED_50_)), giving parameters ED_50_ = 48 mg/kg, bottom = 29.0 °C, and top = 36.1 °C, with R^2^ = 0.83 and DF = 47. The fitted curve and its 95% confidence interval are in black.

**Supplemental Figure 5.**
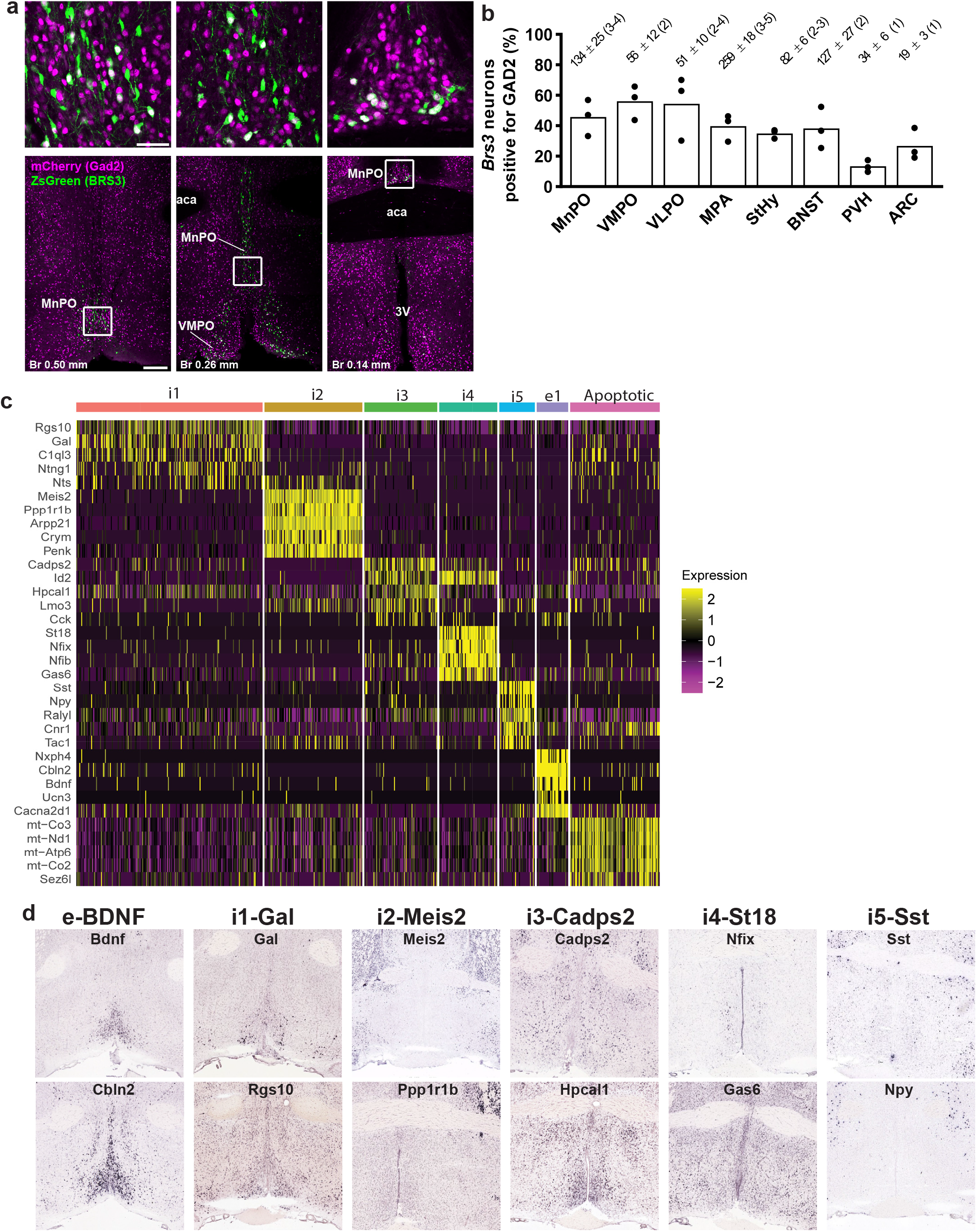
POA^BRS3^ neurons are a mix of excitatory and inhibitory clusters. Related to Figure 5. a) BRS3 (green) and GAD2 (magenta) are expressed in overlapping populations in the MnPO and VMPO in BRS3-Cre;Ai6;Gad2-mCherry mice. Scale bar overview image is 200 µm, inset 50 µm. b) Quantification of BRS3 neurons expressing GAD2. The number of double positive neurons ± s.e.m. in the indicated region and (number of slices/mouse counted) are indicated; n = 3 mice. c) Expression profile of several marker mRNAs for the POA region BRS3 clusters. Data from Moffit et al., 2018. d) Allen Brain Atlas ISH images (http://mouse.brain-map.org/) for two of the marker mRNAs (aside of BRS3 and Vglut2 or Vgat) for each cluster for the POA region.

**Supplemental Figure 6.**
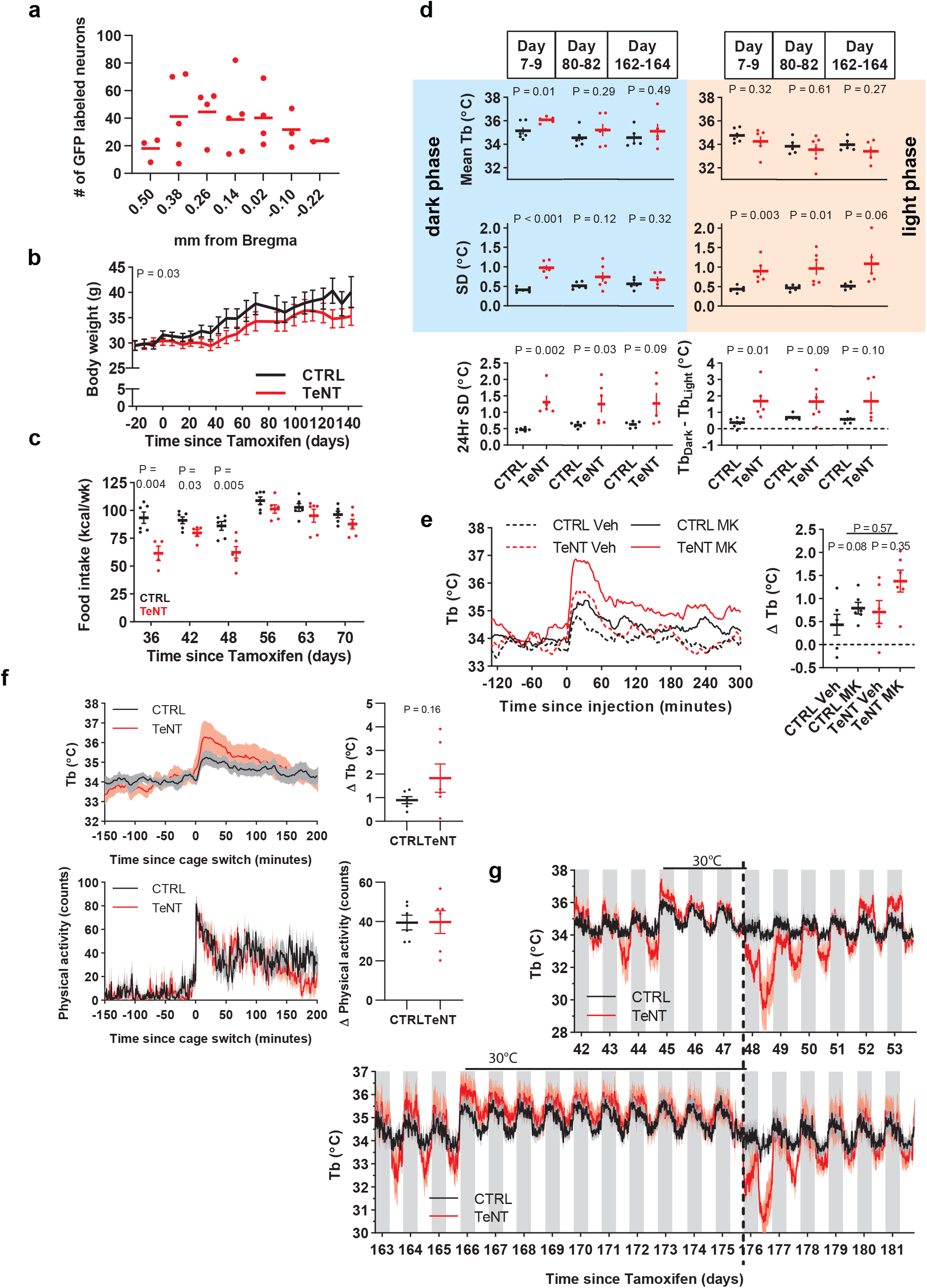
Silencing POA^BRS3^ neurons increases Tb variability and exaggerates Tb changes. Related to Figure 7. a) Number of EYFP labeled (TeNT) neurons in the POA at the indicated distance from Bregma. Every third section was counted from each mouse. b) POA^BRS3^::TeNT mice gain less body weight than do controls. P value, repeated measures ANOVA. c) Reduced food intake in POA^BRS3^::TeNT mice at 4-6 weeks after starting tamoxifen treatment. P values, unpaired t test. d) At indicated number of days after starting tamoxifen treatment, Tb was measured each minute during 72 h intervals and the Tb and standard deviation (SD) during dark (left) and light (right) phases was measured. The circadian amplitude (Tb_dark_-Tb_light_) and the Tb span (95^th^ – 5^th^ Tb percentiles) of the full intervals were also calculated. Data are mean ± s.e.m. P values, unpaired t test. e) Tb response to MK-5046 (10 mg/kg, i.p.) or vehicle (saline) and ΔTb (Tb_60to180_ minus Tb_-150to-30_). Data are mean ± s.e.m. (s.e.m. omitted from left for visual clarity); P value, paired t test between vehicle and MK-5046 and unpaired t test with unequal variance between delta Tb (MK-5046 minus vehicle) of CTRL and TeNT groups. f) Tb and physical activity response to switching mice to a clean cage. ΔTb is Tb_0to60_ minus Tb_- 90to-30_; Δ Physical activity was calculated the same way. Data are mean ± s.e.m. g) Acclimation and Tb response in mice exposed for 3 days (top) vs 10 days (bottom) to 30 °C, otherwise at 22 °C. Data are mean ± s.e.m. In all panels, n=6 mice/group, except for a) with n=5 mice.

**Supplemental Figure 7.**
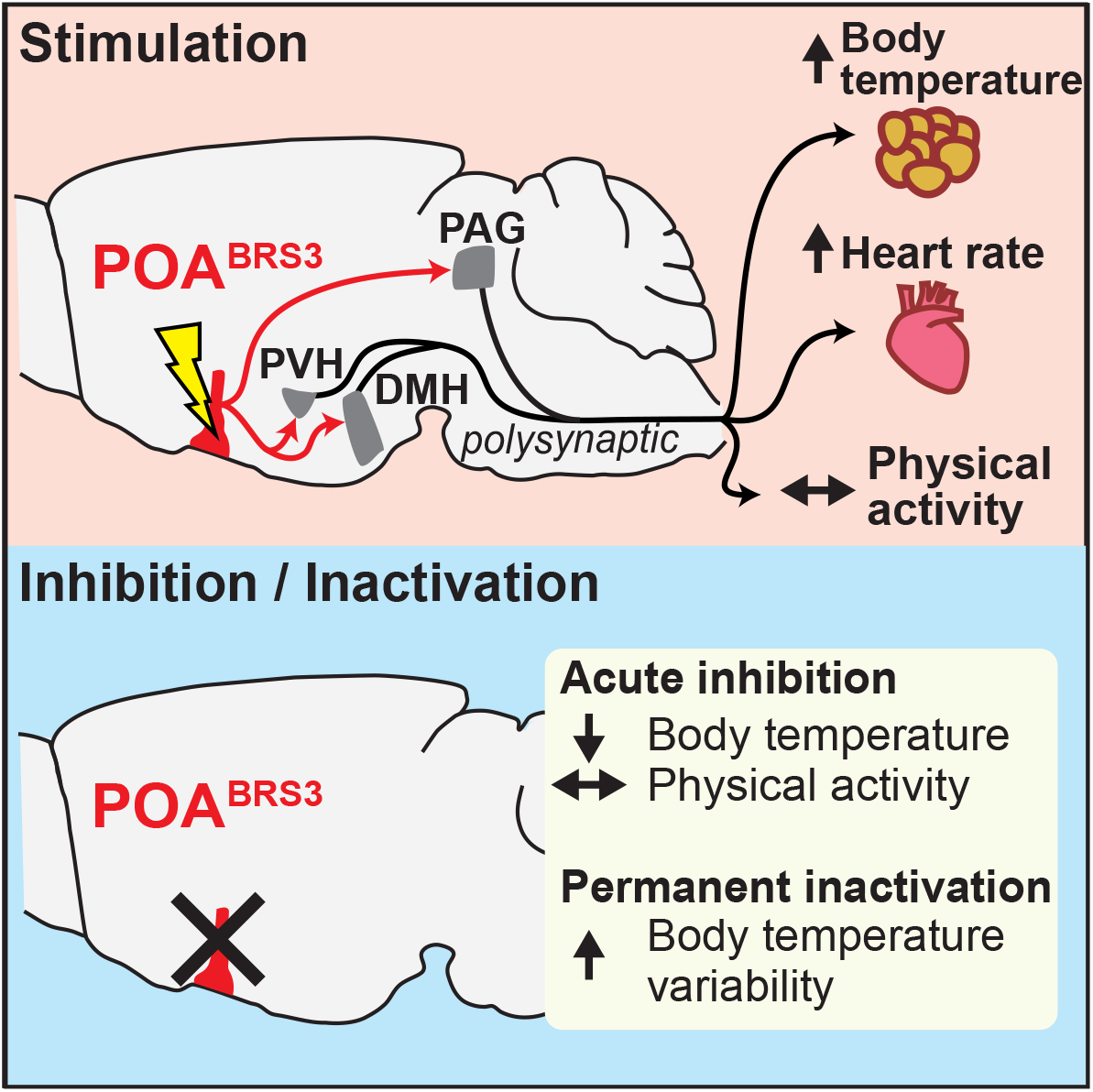
POA^BRS3^ neurons (red) use projections to PVH and DMH to drive sympathetic output to BAT and heart, thereby increasing Tb and HR. POA^BRS3^ neurons also increase Tb through a projection to PAG. Acute inhibition decreases Tb, indicating a roll in cold-defense for POA^BRS3^ neurons. Permanent inactivation increases Tb variability and causes undershoot and overshoot of Tb setpoint during metabolic interventions.

